# Evidence accumulation rate moderates the relationship between enriched environment exposure and age-related response speed declines

**DOI:** 10.1101/2021.10.28.466233

**Authors:** Méadhbh B. Brosnan, Megan H. O’Neill, Gerard M. Loughnane, Daniel J. Pearce, Bryce Fleming, Shou-Han Zhou, Trevor T.-J. Chong, Anna C. Nobre, Redmond G. O Connell, Mark A. Bellgrove

## Abstract

Older adults exposed to enriched environments (EE) maintain relatively higher levels of cognitive function, even in the face of compromised markers of brain health. Response speed (RS) is often used as a simple proxy to measure the preservation of global cognitive function in older adults. However, it is unknown which specific selection, decision, and/or motor processes provide the most specific indices of neurocognitive health. Here, using a simple decision task with electroencephalography (EEG), we found that the efficiency with which an individual accumulates sensory evidence was a critical determinant of the extent to which RS was preserved in older adults. Moreover, the mitigating influence of EE on age-related RS declines was most pronounced when evidence accumulation rates were shallowest. These results suggest that the phenomenon of cognitive reserve, whereby high EE individuals can better tolerate suboptimal brain health to facilitate the preservation of cognitive function, is not just applicable to neuroanatomical indicators of brain ageing, but can be observed in markers of neurophysiology. Our results suggest that EEG metrics of evidence accumulation may index neurocognitive vulnerability of the ageing brain.

**SIGNIFICANCE STATEMENT:** Response speed in older adults is closely linked with trajectories of cognitive ageing. Here, by recording brain activity while individuals perform a simple computer task, we identify a neural metric which is a critical determinant of response speed. Older adults exposed to greater cognitive and social stimulation throughout a lifetime could maintain faster responding, even when this neural metric was impaired. This work suggests EEG is a useful technique for interrogating how a lifetime of stimulation benefits brain health in ageing.

## INTRODUCTION

Cognitive deficits occurring with healthy or pathological ageing catalyse a broad range of challenging consequences (Ball et al., 2010; Barker-Collo and Feigin, 2006; O’Halloran et al., 2013; Prince et al., 2015; Weaver et al., 2009) and are marked by large inter-individual variability (Habib et al., 2007; Norton et al., 2014; Rapp and Amaral, 1992). Robust evidence has emerged over the past three decades demonstrating a powerful positive influence of enriched environments (EE), such as education, leisure and work activities, on the preservation of cognitive function (Cabeza et al., 2019, 2018; Opdebeeck et al., 2016; Stern et al., 2020, 2019a, 1992; Valenzuela and Sachdev, 2006). It has become increasingly apparent that exposure to EE is associated with high levels of cognitive function in older adults, despite structural changes indicative of compromised brain health, a phenomenon commonly referred to as cognitive reserve (e.g. Chan et al., 2018; Stern et al., 1992; Xu et al., 2019). Whether the benefits of EE can compensate for suboptimal neural function revealed by neurophysiological markers remains unexplored.

The speed with which older adults respond to sensory input – hereafter referred to as response speed – has been accepted as a robust index of an individual’s vulnerability to cognitive decline (Bublak et al., 2011; Deary et al., 2010; Gregory et al., 2008; Kochan et al., 2016; Ritchie et al., 2014; Salthouse, 1996). Yet response speed is the aggregate outcome of target-selection, decisional, and motoric computations. Thus, it remains unclear which of these neural processes account for the close association between response speed and neurocognitive health in older adults. Sequential sampling models (including the drift-diffusion model) have offered several explanatory accounts of age-related response slowing (e.g. McKoon and Ratcliff, 2012; Ratcliff et al., 2006a, 2006b, 2004; Ratcliff and McKoon, 2015). However, these models cannot isolate the precise neurophysiological processes driving behaviour. Therefore, the neural mechanisms underpinning age-related declines in response speed remain unclear. To address this question, we have developed EEG tasks and analysis methods that give insight into the underlying selection (early target selection (Loughnane et al., 2016; Zhou et al., 2021), decisional (sensory evidence accumulation (Kelly and O’Connell, 2013; O’Connell et al., 2012; Kelly et al., 2021; Steinemann et al., 2018), and motoric (motor preparatory activity (Kelly et al., 2021; McGovern et al., 2018; Steinemann et al., 2018)) computations that underpin inter-individual differences in response speed (Brosnan et al., 2020; Newman et al., 2017).

A critical extracranial human EEG signal emerging from these investigations is the centro-parietal positivity (CPP). This exhibits the key characteristics of evidence-accumulation signals observed using invasive electrophysiological recordings in animals (Kelly and O’Connell, 2013; O’Connell et al., 2012) and conforms to the dynamics predicted by sequential sampling models in two-alternative choice scenarios (e.g. Kelly et al., 2021; Twomey et al., 2015). In younger adults, the CPP has been repeatedly shown to capture individual variability in response speed (e.g. Brosnan et al., 2020; Murphy et al., 2015; O’Connell et al., 2012). In older adults, recent work on a perceptual decision-making (choice reaction time) task showed that CPP build-up rates were shallower than in a younger control group, indicative of less efficient evidence accumulation (McGovern et al., 2018). However, the specific potential for CPP build-up rate to account for individual differences in response times in older adults, over and above sensory and motoric processes, remains unclear.

The aims of our study were twofold. First, using our EEG framework we tested the hypothesis that neural markers of sensory evidence accumulation (build-up rates of the CPP) would best capture individual variations in speeded target detections, over and above any influence of other neurophysiological processes contributing to the timing of response. Second, we predicted that EE would moderate the relationship between our EEG metrics of evidence accumulation and behaviour. Specifically, that relatively faster response speed would be facilitated by EE, even when the capacity to accumulate sensory evidence, as measured with EEG, was compromised.

## METHODS

### EXPERIMENTAL DESIGN AND STATISTICAL ANALYSES

Seventy-eight healthy volunteers were recruited for this study. Two older adults were excluded due to age ranges more than two standard deviations from the mean (these participants were originally recruited as age-matched controls for a parallel stroke study). A further four older participants were excluded from analysis for various reasons: one was ambidextrous, one was experiencing a current depressive episode and two had scores of 19 and 21, respectively, on the Montreal Cognitive Assessment (MoCA (Nasreddine et al., 2005)), suggesting possible cognitive impairment. The final sample included 31 and 41 older participants (see Table 1 for demographic information). All participants reported being right-handed, had normal or corrected to normal vision, had no history of neurological or psychiatric disorder, and had no head injury resulting in loss of consciousness. Ethical approval was obtained from the Monash Health and Monash University Human Research Ethics Committee prior to the commencement of the study. The experimental protocol was approved and carried out in accordance with the approved guidelines. All participants were volunteers naive to the experimental hypothesis being tested and each provided written informed consent.

**Table 1.** Demographic Information Reported values are M (SD)

|  | Age (yrs) | Gender | Education (yrs) | MoCA |
| --- | --- | --- | --- | --- |
| **Experiment 1** |  |  |  |  |
| Younger Adults<br>($N=31$) | 23.65 (2.87) | 17 female (54.80%) | 15.90 (2.27) | $NA$ |
| Older Adults<br>($N=41$) | 72.41 (5.61) | 26 female (63.40%) | 16.49 (3.48) | 27.46 (1.75) |

### METHOD DETAILS

#### Neurophysiological investigation of response speed

Electroencephalography (EEG) was recorded continuously while participants performed a variant of the random-dot motion perceptual decision-making task (Fig. 1(Newsome et al., 1989; Kelly and O’Connell, 2013; Loughnane et al., 2016; Newman et al., 2017)) During this task, participants fixated centrally and monitored two patches of 150 moving dots (each dot = 6x6 pixels), presented peripherally in each hemifield. During random motion, these dots were placed randomly throughout the patch on each frame. During coherent motion, within one hemifield a proportion (90%) of the dots was randomly selected on each frame to be displaced in either a downward or upward direction on the following frame, with a motion speed of 5° per second. Targets were defined by this seamless transition from random motion to coherent motion (Fig. 1; please note, images in figures 1, 2, and 3 are composite images). Participants signalled target detection with a speeded button press using their right index finger (RT). Targets were separated by intervals of random motion of 1.8, 2.8, or 3.8 s (randomized throughout each block). Targets remained on the screen for 3s, or until the participant pressed the button indicating their detection of coherent motion. The 12 possible trial types (each a combination of one of the 3 periods of random motion, 2 target locations, and 2 coherent motion directions) occurred in a pseudorandom order with the constraint that each different trial type arose twice every 24 trials. All younger adults (*N*=31) performed 8-9 blocks of the task. *N*=22 older adults (who were initially recruited to the study) similarly performed 8-9 blocks, while the remaining older adults (who were later recruited to the study, *N*=19) performed 4-5 blocks of the task at 90 % coherent motion, and a further 4-5 blocks of the task at 25% coherent motion, the latter of which was not analysed for the current study.

**Figure 1.**
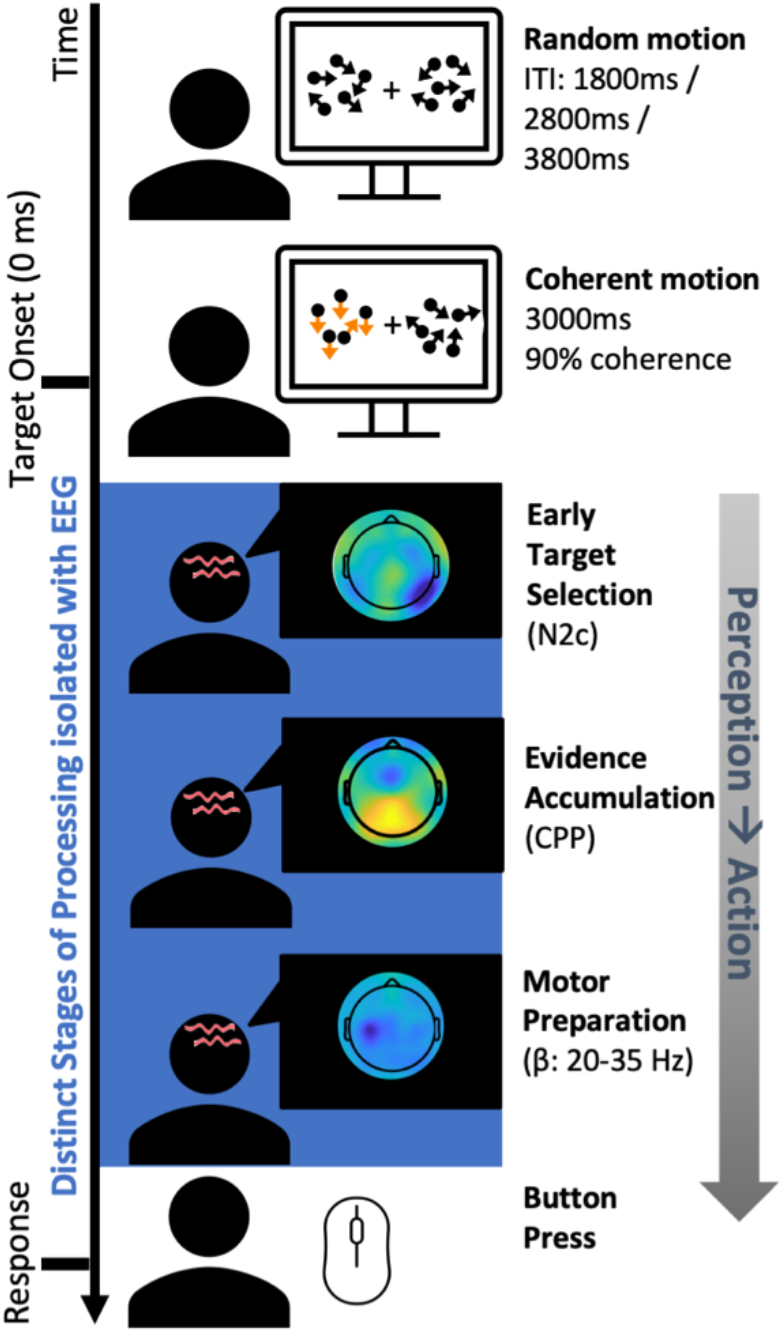
Depiction of the measures obtained on a trial-by-trial basis during the random-dot motion detection task. During the random-dot motion detection task participants fixated centrally while patches of randomly moving dots were presented peripherally (centred at 10° of visual angle either side and 4° visual angle below the fixation square) in both hemifields. During ‘target’ trials, 90% of the dots in one hemifield transitioned from random to coherent motion in either an upward or a downward direction. Targets remained on the screen for 3 s or until the participant pressed the button signalling the detection of coherent motion in either direction. If a fixation break occurred during a trial (either a blink or a gaze deviation >4° left or right of centre), the task halted (stationary dots) until fixation returned to the central fixation dot. Participant response speed was assessed via a right-hand button press for target detection (coherent motion in either upward or downward direction). The blue section illustrates the isolated EEG processes which cannot be obtained from behavioural estimates of speed alone. Each of these processes are derived at each individual trial and collapsed across trials to give an estimation of an individuals’ capacity for each process. Note. ITI denotes inter-target interval.

Critically, a series of *t* tests revealed there were no significant behavioural differences between the older participants recruited for the longer versus shorter task duration(RT *F*_1,.39_ =1.72, *p*=.19; Accuracy *F*_1,39_=.02, *p*=.88) or any of the neurophysiological markers (N2c amplitude *F*_1,39_ =.08, *p*=.77; N2c latency *F*_1,39_=-.10, *p*=.76; CPP onset *F*_1,39_=.82, *p*=.37; CPP slope *F*_1,39_=.67, *p*=.42; CPP amplitude *F*_1,39_=.11, *p*=.74; LHB slope *F*_1,39_=.90, *p*=.35), LHB amplitude *F*_1,39_=.52, *p*=.48), or LHB latency *F*_1,39_=.0, *p*=.99). As such the data were combined to examine the impact of environmental enrichment on neural and behavioural signatures of response speed. All participants were given a short break of 30-60 s between each block. An SR Research EyeLink eye tracker (Eye- Link version 2.04, SR Research/SMI) recorded eye movements, to ensure that participants maintained fixation. The centre of each random-dot motion patch was at a visual angle 10° either side and 4° below the fixation square; each patch covered 8° visual angle and consisted of 150 6 x 6 pixel white dots. If a fixation break occurred during a trial (either a blink or a gaze deviation >4° left or right of centre, detected via EyeLink1000, SR Research Ltd), the task halted (stationary dots). Once fixation returned to the central fixation dot, the trial restarted. The fixation dot remained on screen throughout the entire task; however, the two peripheral patches were only present when the trial was initiated by the participant’s fixation on the central point. The task was run using MATLAB (MathWorks) and the Psychophysics Toolbox extensions (Brainard, 1997; Pelli, 1997; Cornelissen et al., 2002).

#### EEG pre-processing

Continuous EEG was acquired from 64 scalp electrodes using a BrainAmp DC system (BrainProducts), digitized at 500 Hz. Data were processed using a combination of custom scripts and EEGLAB (Delorme and Makeig, 2004) routines implemented in MATLAB (MathWorks). First, noisy channels were identified using visual inspection of channel variances across the entire recording, to be interpolated at a later stage below. Next, the EEG was detrended, then notch filtered at 50, 100 and 150 Hz to eliminate line noise and its harmonics, then high-pass filtered at 0.1 Hz using a Hamming windowed sinc FIR filter via EEGLAB. Channels with zero or extreme variance identified from the first inspection were interpolated via spherical spline. A 35-Hz low-pass filter was then applied to the data using Hamming windowed sinc FIR filter also, and the data were re-referenced to the average reference. Epochs were extracted from the continuous data from -200 to 1500 ms from target onset. For both the ERP and stimulus-aligned LHB signals, the epochs were baselined with respect to -100 to 0 ms before target onset. For the response-aligned beta waveforms, the data were baselined between -450 to -350ms pre-response. Using triggers recorded by the EEG, we defined trials as the period between the beginning of random (i.e. non-target) motion and either a valid response, a fixation break, or the onset of the next period of random motion (i.e. a non-response). To minimise the interaction between overlapping ERP components, the data were subjected to Current Source Density transformation with a spline flexibility of 4 (Kayser and Tenke, 2006).

A trial was excluded from the analysis if any of the following conditions applied: (1) if RTs were ≤150 ms (pre-emptive responses) or ≥ 1800 ms (responses after coherent motion offset); (2) if the EEG from any channel exceeded 100 µV during the interval from 100 ms before target onset to 100 ms after response; or (3) if central fixation was broken by blinking or eye movement 3° left or right of centre, during the interval between 100 ms before target onset and 100 ms after response. Please note that EyeLink data were not saved for N=5 out of the N=41 older adults due to a technical error and this final step was therefore not included for this subset of participants. Nonetheless fixation was monitored in real-time using EyeLink during task performance as described in the preceding section so no trials with eye movements >4° from centre were included.

With the remaining trials for each participant, CPP and N2 waveforms were aggregated by averaging the baseline-corrected epochs, for right and left hemifield targets at the relevant electrode sites. The N2c component was measured contralateral to the target location, respectively, at electrodes P7 and P8 (Brosnan et al., 2020; Loughnane et al., 2016; Newman et al., 2017) and the CPP was measured centrally at electrode Pz (Kelly and O’Connell, 2013; Loughnane et al., 2016; Newman et al., 2017; O’Connell et al., 2012; Twomey et al., 2015). Subsequently, N2c latency was identified on a subject level as the time point with the most negative amplitude value in the stimulus-locked waveform between 150-400 ms, whereas N2c amplitude was measured as the mean amplitude inside a 100 ms window centred on the stimulus-locked grand average peak of the N2c collapsed across hemifield (Loughnane et al., 2016). Onset latency of the CPP was measured by performing running sample-point-by-sample-point t tests against zero using a 25 ms sliding window across each participant’s stimulus-locked CPP waveforms. CPP onset was defined as the first point at which the amplitude reached significance at the 0.05 level for 90 consecutive points (Foxe and Simpson, 2002; Kelly et al., 2008; Loughnane et al., 2016). CPP build-up rate was defined as the slope of a straight line fitted to the response-locked waveform (Brosnan et al., 2020; Loughnane et al., 2016; O’Connell et al., 2012), with the time window defined individually for each participant from -150 to 50 ms post-response (e.g. Brosnan et al., 2020; Stefanac et al., 2021). CPP amplitude was measured as the mean CPP amplitude between -50 and +50 ms around the participants’ individual response (e.g. Kelly and O’Connell, 2013; Van Kempen et al., 2019).

Finally, LHB power was calculated using the temporal spectral evolution approach (Thut et al., 2006). All epochs were bandpass filtered between 20-35 Hz, converted to absolute values (rectified) and trimmed by 200 ms at each end of the epoch to remove filter warm-up artefacts. The data were then smoothed by averaging within a 100-ms moving window, moving incrementally forward in 50-ms increments. LHB latency was measured within the left hemisphere motor site (C3) (corresponding to the right handed response modality) as the most negative-going point between 0 and 1000 ms. Beta slope was defined as the slope of a straight line fitted to the response-locked waveform, with the time window defined individually for each participant between 300 to 50 ms pre-response. Beta amplitude was measured as the mean amplitude of a 100 ms window centred on a participants’ response (i.e., -50 to +50 ms around response).

#### Drift diffusion modelling

A hierarchical drift-diffusion model was used to analyse behaviour in the older and younger adult groups (Wiecki et al., 2013; Gee et al., 2018). The RT distributions for the response and no-response choices were first split into five bins according to the quantiles 0.1, 0.3, 0.5, 0.7 and 0.9. Together with a single bin containing the number of missed responses, these six bins (five response bins and one missed responses bin) were then used to fit a drift-diffusion model using the G-square method (Ratcliffe et al., 2016; Gee et al., 2018). The proportion of responses within each bin was determined by subtracting the cumulative probabilities for each successive quantile from the next highest quantile. These proportions were then multiplied by the number of observations to obtain the expected frequencies *E* ∈ *R*. Next, the observed proportions, determined from the data (in this case: 0.1, 0.2, 0.2, 0.2, 0.2 and 0.1), were also multiplied by the number of observations to obtain the observed frequencies *O* ∈ *R*. G-square was then calculated as

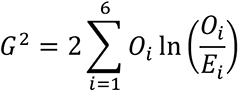

where *i* ∈ *N* represents the quantile number. The drift-diffusion model parameters *α*, *v* and *t* were then determined by minimising the G-square statistic using the modified Powell method (Powell, 1964). The fitted drift-diffusion model assumed that the response caution, mean drift rate across trials, and non-decision time varied with the response bins. Further details can be found in Ratcliffe et al. (2018).

#### Assessment of Environmental Enrichment

Participants completed the Cognitive Reserve Index questionnaire (CRIq) (Nucci et al., 2012), a standardised semi-structured interview designed to estimate an individual’s level of lifetime cognitive enrichment through a formal computational model. This model encompasses an individual’s education, work and leisure activities across the lifespan with consideration given to the participant’s age, providing both an overall age-stratified and standardised Cognitive Reserve Index (CRI) and individual standardised subscale scores for each of the three components. One participant did not complete the CRIq due to time constraints. For this participant, we imputed their scores on all four CRI measures using the mean from the rest of the sample.

Participants first reported the number of years in which they had engaged in formal education and additional vocational training. All occupations held since the individual was 18 years old were categorised using the five-point point scale provided by the CRI. These ranged from low skilled manual work (e.g. level 1 includes occupations like call centre operator, and gardener) to highly responsible or intellectual occupation (e.g. level 5 includes managing director of a big company or surgeon). Participants were additionally asked about their involvement in leisure activities that may be repeated with varying frequencies over the lifetime, including but not limited to reading, volunteering, socialising, managing accounts, going on holidays/trips. Activities were grouped into weekly, monthly, annual and fixed frequency activities, and then into whether they were completed never, rarely, often or always, and for how many years of life. Participant engagement in each of these domains is summarised in Supplementary Tables 10-12.

#### Data processing

Outliers were defined in SPSS using the interquartile range (IQR), separately for the younger and older adults. The interquartile range is the 3rd quartile (75th percentile) minus the 1st quartile (25th percentile). A value was identified as an outlier if either of the following conditions were met: if the value was <25th percentile - 3*IQR or if the value was >75th percentile + 3*IQR. Using this method, no outliers were identified on any of the behavioural, EEG, or EE measures used in the analyses below.

#### Code Availability

Our internally developed EEG pipeline including the preprocessing steps and isolation of the EEG metrics can be found here https://github.com/gerontium/big_dots, openly available under a Creative Commons Attribution-NonCommericial-ShareAlike 3.0 International License.

The code for the analysis exploring the minimum trial numbers needed to reliably isolate the CPP build-up rate as an EEG marker of sensory evidence accumulation rate can be found on OSF at the following link: https://osf.io/vupa4/files/osfstorage/640391f8c7472300bb10aeea, openly available under a.CC-By Attribution 4.0 International License

The code for the hDDM of the behavioural data can be found here: https://github.com/shou-han/DetectionDDM.

Note that our data were collected under a larger, multi-center international study using EEG to gain mechanistic insights into perceptual decision-making deficits occurring post stroke. Our ethics does not permit us to share data from the project.

### STATISTICAL ANALYSIS

#### The Relationship between Age, Behaviour and EEG

To assess age-related differences in behaviour, two one-way ANOVAs were conducted on Accuracy and RT. Next, to test whether the older and younger adults differed across N2c, CPP, and LHB dynamics, eight one-way ANOVAs were conducted with the EEG variables (N2c latency, and amplitude, CPP onset, build-up rate, and amplitude, LHB build-up rate, LHB amplitude, and LHB latency) as dependent variables, and age as a factor. To assess whether inter-individual differences in RT on the perceptual decision-making paradigm (RT) varied as a function of EEG signals of perceptual decision-making, the EEG parameters which differed in older versus younger adults (BF_10_>1) were each added sequentially into regression models in a hierarchical fashion(Newman et al., 2017). Order of entry was determined by the temporal order in the perceptual decision-making process: early target selection (N2c latency); evidence accumulation (CPP onset, build-up rate, and amplitude), and motor preparation (LHB build-up rate, LHB latency, and LHB amplitude). This hierarchical entry method was implemented to assess whether each of the separate neurophysiological signals improved the model fit for RT over and above the signals that temporally preceded them. All neurophysiological signals that improved the model fit for RT were entered into a separate regression model to obtain accurate parameter estimates. Age was entered as the first predictor, centred (i.e., all raw scores for each participant were subtracted from the mean score of the variable) to reduce multicollinearity. Please note all statistical tests were two-sided. Effect sizes of regression models were calculated using Cohen’s *F*^2^ using the following formula: (*R*^2^/(1-*R*^2^)). Behavioural data was visualised using *RainCloudPlots* ^f^or MATLAB (Allen et al., 2018, 2019). The EEG signals were visualised using GRAMM for MATLAB (Morel, 2018).

#### Moderation Models

To elucidate the moderating effects of evidence accumulation rate, amplitude, and beta latency on the relationship between EE and response speed, three moderation analyses were performed using the PROCESS computational toolbox (Hayes, 2012, 2014), *Bonferroni*- corrected for multiple comparisons (alpha .05/3 moderation models).

#### Confirmatory Bayesian Analyses

For the confirmatory Bayesian modelling, results were compared with the null model, and JASP default settings were used (for the regression and ANOVA analyses: JZS prior (JZS (Rouder and Morey, 2009); regression analyses: r scale .354, ANOVA analyses: r scale fixed effects .5; Cauchy prior of .707 for the one sample t-test). BF_inclusion_, or BF10 values are reported throughout and can be interpreted such that values above 1 indicate strength of evidence in favour of the alternative and values below 1 strength of evidence in favour of the null hypothesis.

#### Minimum Trial Analysis

The minimum trial analyses included all participants (N=53) who completed 8 or more blocks of the task. One individual was identified as an outlier (>2SD from the mean) with regards the number of trials included (N=109 trials) and was excluded, therefore resulting in a total of *N*=52 participants. All of these 52 participants had a minimum of 129 valid response-locked trials, which we used to investigate remaining questions (*M*=183.06; *SD*=19.95; range 129-207).

We first created new estimates of both RT and CPP build-up rate by randomly selecting *N* trials (either 20, 40, 60, 80, 100, or 120) from the total pool of 129 trials. We repeated this random data-sampling using *N* trials, 1000 times for each bin size. Accordingly, for each participant, we derived 1000 estimates of RT and CPP build-up rate for each of the six trial sizes. We then tested whether the likelihood that the estimates of RT and CPP build-up rate were more likely to deviate from the *true* mean estimates with reduced trial numbers. We addressed this question using two approaches. First, we calculated the signal to noise ratio (SNR) of the CPP build-up rate and RT (calculated as mean / standard deviation) and ran two repeated measures ANOVAs (again with trial bin as the repeated measure). Next, to verify this pattern of results, we ran Kolmogorov-Smirnov tests on the mean estimates of CPP build-up rate and RT to assess whether the cumulative distribution function (CDF) increased with each reduction in trial number.

In the results reported in the main body of the text, we demonstrated a large effect size for the relationship between CPP build-up rate and RT (Pearson’s *r*)= -.60. Cohen’s (1988) cut-off for a large effect size is .5. As such, we defined the minimum number of trials at which a reliable CPP estimate can be derived as the number at which we can observe a strong effect size (i.e., an effect size greater or equal to .5) for the relationship between RT and CPP build- up rate. To investigate this, we calculated the direct relationship, using Pearson’s correlation, between CPP build-up rate and RT for each of the 1000 permutations for each of the 6 bin sizes (20 up until 120 trials) using MATLAB. We then ran a Bayesian one sample *t*-test to test whether the estimates of effect size (*r*) for each bin size were significantly larger than -.5 using JASP. For this minimum trial Bayes Factor analyses the alternative hypothesis was set at measure 1 ∼= measure 2.

### The Relationship between hDDM parameters and EEG markers of decision making

To examine the association between the hDDM parameters derived using computational modelling and the neurophysiological signals of decision making derived using EEG, we modelled each hDDM parameter (non-decision time *t*, drift rate *v*, and response caution *a*) as a function of the EEG signals. Specifically, we used three hierarchical linear regression models to model the three hDDM parameters as a function of the neural metrics, namely early target selection (N2c amplitude, N2c latency), sensory evidence accumulation (CPP onset, CPP build-up rate, CPP amplitude) and motor preparation (LHB latency). The EEG signals were entered hierarchically into the models based on the temporal order in which they occur (as per the modelling procedure for response speed described in detail above). Age was entered as a nuisance variable at the first stage of each model (centred to avoid multicollinearity).

#### The Influence of Age on the hDDM parameters

To explore whether older adults differed from their younger counterparts on the three hDDM parameters (non-decision time *t*, drift rate *v*, and response caution *a*), three independent t-tests were run using a *Bonferroni*-correction for three multiple comparisons (alpha corrected threshold of p<.02).

## RESULTS

### Individuated neurophysiological metrics indexing visual response speed were isolated using a perceptual-decision making EEG task

Seventy two participants (41 older *M*=73 years, *SD*=5, range = 63-87 and *N*=31 younger *M*=24 years, *SD*=3, range = 18-28) performed a variant of the random-dot motion task (Newsome et al., 1989) while 64-channel EEG was recorded to isolate neurophysiological processes along the perception to action continuum. We have developed this formal framework for parsing discrete EEG metrics (Newman et al., 2017; Brosnan et al., 2020) to estimate an individuals’ capacity for a given neurophysiological processes (Fig. 1). For instance, we have recently demonstrated the utility of this framework for linking individual differences in MR markers of structural and functional connectivity, neurophysiology, and behaviour in younger individuals (Brosnan et al., 2020). In the current study, we employ the same approach to isolate eight distinct and previously validated neural metrics (Newman et al., 2017; Brosnan et al., 2020), namely early target selection (N2c amplitude and latency (Loughnane et al., 2016)), sensory evidence accumulation (CPP starting point (onset latency), build-up rate (slope), and decision bound (amplitude) (O’Connell et al., 2012; McGovern et al., 2018; Steinemann et al., 2018), and motor preparation (left hemisphere beta (LHB) build- up rate (slope), timing (stimulus-aligned peak latency), and threshold (amplitude (O’Connell et al., 2012; McGovern et al., 2018) see Fig. 1.

### Response Speed Measures Are Sensitive to both Age and EE

During the simple random-dot motion detection task, participants fixated centrally while two patches of randomly moving dots were presented to the periphery (see Fig. 1). A target was defined as 90% of the dots in one hemifield transitioning from random to coherent motion, in either an upward or downward direction. Participants were required to respond to any coherent motion (i.e., in either direction) with a right-handed button press. Behavioural analyses (Fig. 2A) indicated that this task was sensitive to age-related deficits in response speed. The older adults were markedly slower at responding, as evidenced by significantly slower response times (RTs) to the visual targets relative to the younger adults (*F*_1,70_ =38.34, *p*<.001, partial *η*^2^= 0.35, BF_10_=287907.20; older *M*=593.33ms, *SD*=125.40; younger *M*=439.05ms, *SD*=67.87; Fig. 2A). Target detection accuracy was high for the overall sample 95.92% (*SD*=5.29, range 71-100%), but nonetheless the older adults were less accurate at detecting coherent motion than their younger peers (older: *M =* 94.40%, *SD* = 6.3%; younger: *M*=97.90, *SD*=2.60%; *F*_1,70_ = 8.12, *p*=.006, partial *η*^2^ = 0.10, BF_10_=7.17). Critically, the age-related declines in RTs remained significant even after covarying for differences in accuracy (*F*_2,69_ = 27.16, *p*<.001, partial *η*^2^= 0.44).

**Figure 2.**
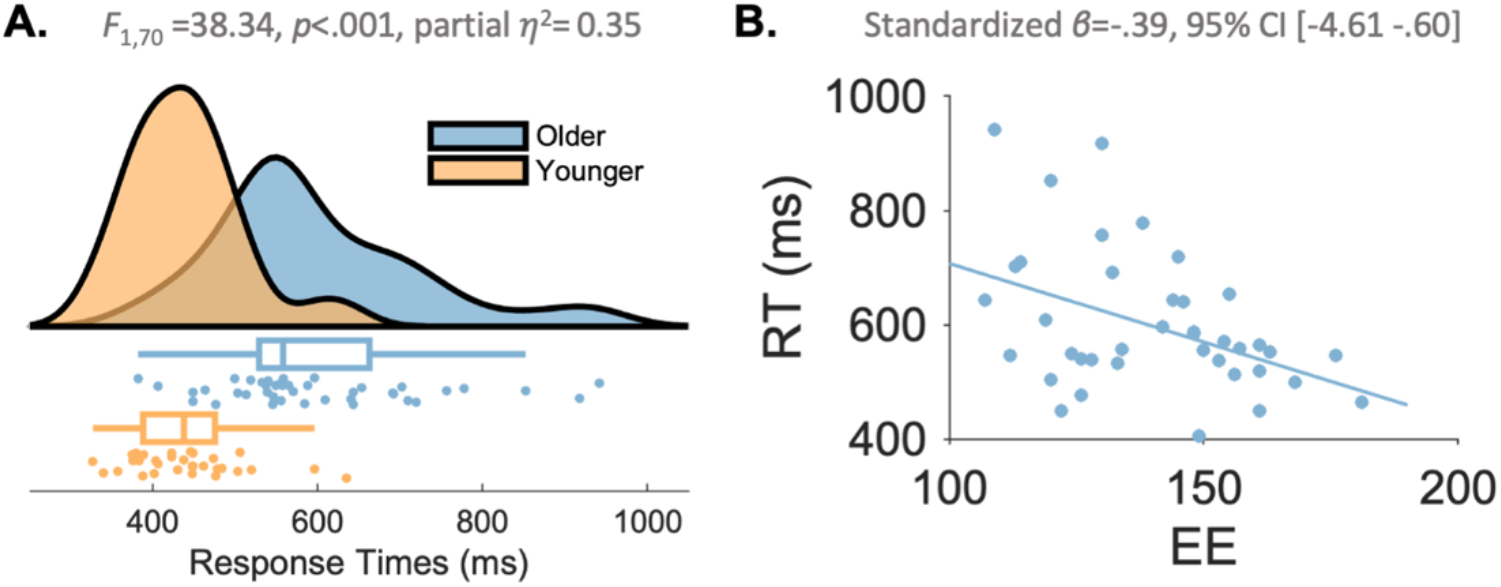
Response Speed Measures on the Decision Task Sensitive to both Age and EE. **A.** Healthy ageing was associated with markedly slower response times (RT) to perceptual targets, with large interindividual differences in response speed. During a variant of the random-dot motion task older participants were, in general, slower to respond relative to their younger peers, suggesting this measure was sensitive to age-related deficits in response speed. Each individual dot represents a participant (lower panel) and the upper/lower edges of the whiskers represent the upper/lower quartiles plus/minus 1.5 × the interquartile range. The distribution is captured by a violin plot for the two groups (upper panel). *Note*. **B.** A lifetime of enriched environments (EE), captured by the composite score of Cognitive Reserve Index Questionnaire (CRIq; Nucci et al., 2012) varied according to individual differences in response speed in the older adults. This effect was driven by the Leisure subscale of the assessment which is visualised here as a function of RT.

We next sought to verify previously reported associations between a lifetime of EE and response speed e.g. (Lee et al., 2014; Park et al., 2014). For this, we modelled RT from the random-dot motion task as a function of environmental enrichment using the Cognitive Reserve Index questionnaire (CRIq (Nucci et al., 2012)) in the older adult cohort only. The CRIq is a previously validated semi-structured interview which assays levels of cognitive stimulation through the assessment of three domains of activity throughout an individual’s lifetime: *Education, Work Activities,* and *Leisure Activities* (see methods for details). As the neuroprotective effects of EE are posited to accumulate over the course of a lifetime (Robertson, 2014), we collected this information in the older cohort only.

As expected, this model was statistically significant, and EE (the overall model) explained 20.5% of the variance of RT (*R*^2^_adj_ =.21*, F_3_*_,36_=4.36, *p*=.01, partial η^2^=.27). Consistent with previous work (Lee et al., 2014; Park et al., 2014), this effect was driven by the CRI *Leisure* subscale, which accounted for independent variance in the modelling of RT (Standardized *β*=-.45, *t*=-3.13, *p*=.003, 95% CI [-4.98 -1.06]), such that older adults with greater exposure to enriched leisure activities exhibited faster visual response speeds (Fig. 2B). In contrast, neither CRI *Education* (Standardized *β*=.06, *t*=.41, *p*=.69, 95% CI [-2.73 4.10]) nor CRI *Work* (Standardized *β*=.31, *t*=1.94, *p*=.06, 95% CI [-.09 3.93]) accounted for independent variance in RT. In order to obtain accurate parameter estimates for the relationship between CRI *Leisure* and RT, not influenced by the non-informative signals, CRI *Leisure* was entered into a separate linear regression model. This model explained 13.2% of the variance (Cohen’s *F*^2^=.18) in RT (Standardized *β*=-.39, *t*=-2.63; *F*_(1,38)_=6.93, *p*=.01, 95% CI [-4.61 -.60], partial η^2^=.15, Fig. 2A).

Bayesian Linear Regression analyses modelling RT as a function of each CRI subscale provided additional support for the results of the frequentist statistics. Any model including CRI *Leisure* indicated a Bayes Factor at least 2.9 times more in favour of H_1_ than H_0_ (Extended Data Fig. 2-1). In contrast, Bayes Factors for both CRI *Work* and CRI *Education* (independently and combined) provided anecdotal to very strong evidence for the null hypothesis (i.e., there was no evidence to suggest that these factors account for independent variance in RT; all three BF_10_<.88 and >.03). This suggests that an individual’s leisure engagements help to mitigate age-related declines in visual response speed. An exploratory analysis conducted to investigate which specific aspects of leisure activities may have contributed to this effect implicated using modern technology (*t*_39_=-4.37, *p*<.001, BF_10_=240.49), engaging in social activities (*t*_26.88_=-4.49, *p*<.001, BF_10_=106.02), attending events such as conferences, exhibitions and concerts (*t*_32_=-3.98, *p*<.001, BF_10_=10658.60), and vacationing (*t*_24.83_=3.11, *p*=.01, (BF_10_=72.46; see Extended Data Fig. 2-2- 2-5 for further information).

### Individual differences in response speed are captured by neural metrics of sensory evidence accumulation rate

The analyses thus far have confirmed age-related differences in behavioural markers of response speed - a validated behavioural measure of cognitive resilience. We next sought to understand how each neural metric related to individual differences in response speed using a hierarchical regression model to isolate the contribution of each neural metric, over and above those which temporally preceded it.

To determine the explanatory power of the neurophysiological signals for predicting behaviour**, age** was entered as a nuisance variable (centred to avoid multicollinearity) in the first step of the model. Unsurprisingly, this offered a significant improvement in model fit, as compared with the intercept-only model (*R*^2^_adj_ =.36, *p*<.0005 Fig. 3). Neither the marker of early target selection (N2c latency (*R*^2^_adj_ = .37, *p*=.21), N2c amplitude (*R*^2^_adj_ =.37, *p*=.22) nor the starting point of the evidence accumulation process (CPP onset; *R*^2^_adj_ =.37, *p*=.58) offered any additional improvement in model fit (Fig. 3A).

**Figure 3.**
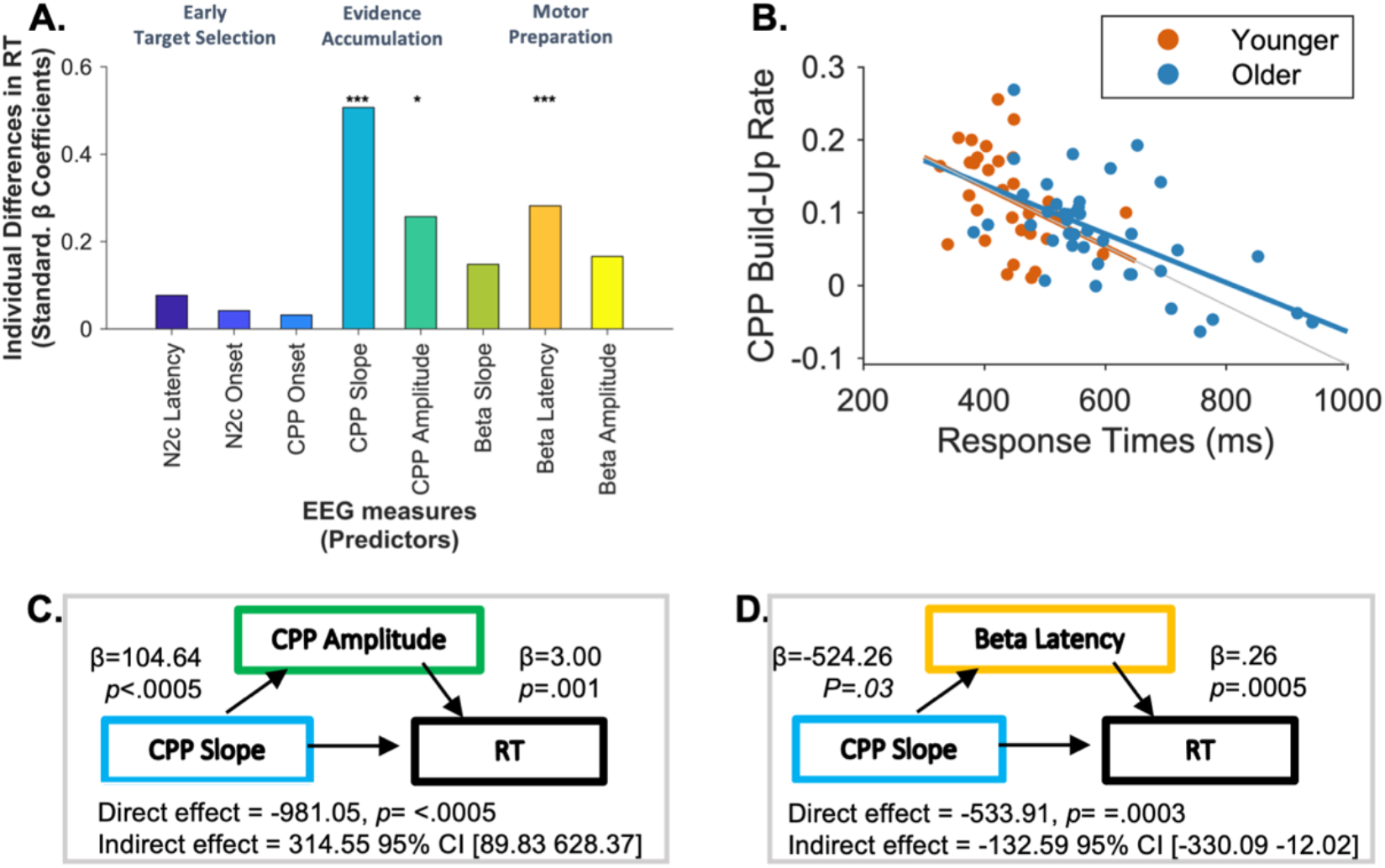
Individual differences in response speed are captured by sensory evidence accumulation build-up rate. **A.** Associations between response speed (RT) and the EEG variables. Results from the final regression model of RT are reported in Table 1. *Note* the absolute value of all standardised beta values were plotted for visualisation purposes and nuisance variables entered from the first step in the model are not visualised here. **B.** The relationship between CPP build-up rate and RT for older and younger adults. CPP build-up rate was directly associated with an individuals’ response speed. Moreover, an individuals’ capacity to accumulate sensory evidence indirectly impacts response speed by influencing CPP amplitude **(C)** and Beta latency (**D**), both of which mediate the association between CPP build-up rate and response speed. For further information and visualisation of the neural metrics see Extended Data Fig. 3-2, Fig. 3-3, and Fig. 3-4.

Evidence accumulation build-up rate, indexed via the <u>CPP build-up rate</u> significantly improved the model performance, accounting for an additional 17% of the variance (*R*^2^_adj_ =.54, *R*^2^_change_ =.17, *p*<.0005, Fig. 3A, Table 2), such that steeper CPP slopes, indicative of a faster build-up rate of sensory evidence, were associated with faster response speeds (Fig. 3B). Adding <u>CPP amplitude</u> offered a further significant improvement in the model, such that individuals with lower CPP amplitudes showed faster RTs (*R*^2^_adj_ =.61, *R*^2^_change_ =.07, *p*=.001).

**Table 2.**
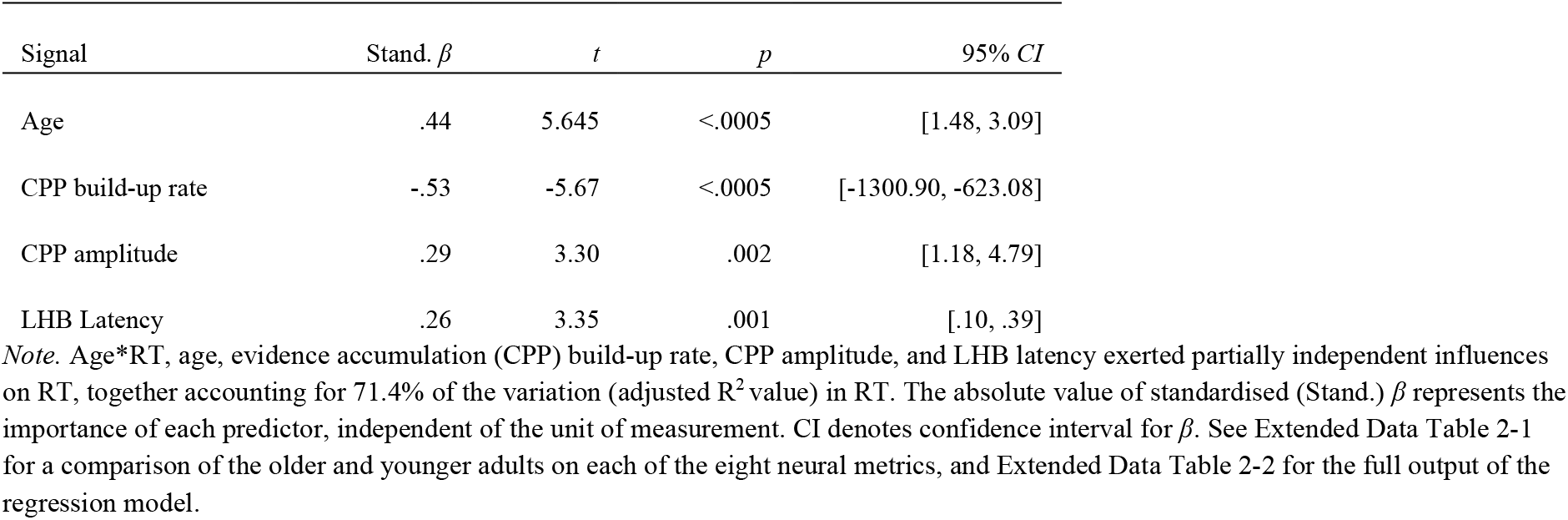
Parameter estimates from the final linear regression model for reaction time (RT) as a function of the neurophysiological signals

| Signal | Stand. $\beta$ | $t$ | $p$ | 95% CI |
| --- | --- | --- | --- | --- |
| Age | .44 | 5.645 | <.0005 | [1.48, 3.09] |
| CPP build-up rate | -.53 | -5.67 | <.0005 | [-1300.90, -623.08] |
| CPP amplitude | .29 | 3.30 | .002 | [1.18, 4.79] |
| LHB Latency | .26 | 3.35 | .001 | [.10, .39] |

*Note.* Age\*RT, age, evidence accumulation (CPP) build-up rate, CPP amplitude, and LHB latency exerted partially independent influences on RT, together accounting for 71.4% of the variation (adjusted $R^2$ value) in RT. The absolute value of standardised (Stand.) $\beta$ represents the importance of each predictor, independent of the unit of measurement. CI denotes confidence interval for $\beta$ . See Extended Data Table 2-1 for a comparison of the older and younger adults on each of the eight neural metrics, and Extended Data Table 2-2 for the full output of the regression model.

While the build-up rate of motor preparation (LHB slope) explained no additional variance in RT (*R*^2^_adj_ =.59, *R*^2^_change_ =0, *p*=.94), <u>stimulus-aligned LHB peak latency</u> significantly improved the fit, such that an earlier peak latency of this motor preparatory marker was associated with faster RT (*R*^2^_adj_ =.65, *R*^2^_change_ =.05, *p*=.003). Finally, adding LHB amplitude offered no significant improvement in the model (*R*^2^_adj_ =.65, *R*^2^_change_=01, *p*=.17). A post-hoc power analysis indicated that with 72 participants, 8 tested predictors (8 neural metrics), age as a control variable, and an effect size Cohen’s f^2^=.29 (based on final regression model), 88.86% power was achieved G*Power 3.1).

In order to isolate the variables explaining independent variance in RT over and above that explained by other non-informative signals, age, CPP build-up rate, CPP amplitude, and LHB peak latency were entered into a single separate linear regression model. When these four independent variables were included in the final model, they <u>accounted for 65.6% of the</u> <u>variation in RT</u> (*F*_4, 66_=34.43, *p*<.0005, Table 2).

Next, to further establish the utility of these signals as specifically sensitive to individual differences in ageing, we repeated this linear regression model (with CPP build-up rate, CPP amplitude, and LHB peak latency), just for the older cohort. This model accounted for 48.7% of the variation in RT (*F*_3, 37_=13.66, *p*<.0005) and CPP build-up rate (stand. β=-.66, *p*<.0005), CPP amplitude (stand. β =.27, *p*=.05) and LHB Latency (stand. β=-.31, *p*=.013) all accounted for independent variance in response speed.

Finally, to validate these results, we ran some confirmatory Bayesian analyses. In keeping with the frequentist analyses, the Bayesian regression model for RT indicated strong support for the alternative hypothesis for age (BF_10_=1028.52), CPP slope (BF_10_=4732.19), CPP amplitude (BF_10_=17.67), and LHB latency (BF_10_=31.74). There was no statistical evidence to suggest that N2c amplitude (BF_10_=.72), N2c latency (BF_10_=.62), CPP onset (BF_10_=.55), LHB build-up rate (BF_10_=.68), or LHB amplitude (BF_10_=.75) influenced RT (see Extended Data Fig. 3-1 for more detailed results from this Bayesian linear model).

Together the findings above indicate that CPP build-up rate, CPP amplitude, and LHB latency exerted direct and partially independent influences over RT. On the basis of previous work, we assume that the impact of both CPP amplitude and LHB latency on RT is, at least in part, determined by accumulated sensory evidence, reflected in the temporally preceding CPP build-up rate (O’Connell et al., 2012; Kelly and O’Connell, 2013; Steinemann et al., 2018; Brosnan et al., 2020). We tested this here by assessing whether the influence of CPP amplitude and LHB latency on RT was mediated by CPP build-up rate. In both cases, bootstrapped mediation analyses (5000 samples) indicated that this was the case (CPP build-up rate→CPP amplitude→RT indirect effect 281.98, bootstrapped SE 168.06, CI [18.00 669.46]; CPP build-up rate→ LHB latency→RT indirect effect -220.68, bootstrapped SE 102.48, CI [-459.99 -58.22; Fig. 3C, D]). This demonstrates that variability in age-related deficits in RT captured by CPP amplitude and LHB latency are dependent, at least partly, on individual differences in the rate at which sensory evidence can be accumulated. These results suggest that the CPP build-up rate constitutes a critical contributor to interindividual differences in response speed.

### Neural metrics of evidence accumulation build-up rate moderate the relationship between environmental enrichment and response speed

The results thus far demonstrate that both levels of environmental enrichment and task-related neural metrics (particularly the build-up rate of evidence accumulation) are strong determinants of age-related individual declines in behaviour (response speed) at an inter-individual level. This raises the possibility that the relationship between EE and response speed might differ according to individual differences in evidence accumulation build-up rate. It is well established within the (neuro)cognitive reserve literature that high EE individuals can preserve relatively high levels of cognitive function, despite suboptimal structural markers of brain health (e.g., grey matter atrophy).

Signals displaying evidence accumulation dynamics are not specific to a single cortical area but rather have been found in a number of regions throughout the brain (e.g., Ratcliff et al., 2003; Cisek and Kalaska, 2005; Huk, 2005; Ding and Gold, 2010; Pape and Siegel, 2016), (e.g., Ratcliff et al., 2003; Cisek and Kalaska, 2005; Huk, 2005; Ding and Gold, 2010; Pape and Siegel, 2016), and therefore may represent a global, widespread neurophysiological process. Previous work investigating cognitive reserve using similarly widespread, global measures of brain structure (e.g. grey matter atrophy; Chan et al., 2018), and neuropathology (e.g., amyloid plaques and tangles in Alzheimer’s patients; Xu et al., 2019) has demonstrated that higher EE individuals can maintain better cognitive performance despite these compromised markers of brain health. Here, we hypothesised that this same phenomenon would be observed using a neurophysiological marker of ageing brain health, i.e., neural metrics of sensory evidence accumulation rate. Specifically, having established evidence accumulation build-up rates as the critical neural marker indicative of the maintenance of response speed, we tested the hypothesis that high EE individuals would show relatively preserved response speeds, even when this core neurophysiological process was impaired.

To address this, we tested whether each of the three neural markers significantly moderated the relationship between CRI leisure and RT using three separate moderation models, *Bonferroni*-corrected for multiple comparisons (alpha .05/3 moderation models => alpha-corrected threshold = .016). These results revealed a specific moderating influence of CPP build-up rate on the association between EE (CRI Leisure) and RT, as evidenced by a CRI Leisure by CPP build up rate interaction (coefficient = 32.00, se=10.55, *t*=3.03, *p*=.005, CI [10.61 53.39]), which remained significant when covarying for (age; coefficient = 31.45, se=10.81, *t*=2.91, *p*=.006, CI [9.51 53.38], Table 3, Fig. 4). In contrast, no moderating influence was observed for CPP amplitude (coefficient = .14, se=09.71, *t*=1.39, *p*=.17, CI [-.06 .33]) or LHB latency (coefficient = -.02, se=.006, *t*=-2.48, *p*=.02, CI [-.03 0]). Follow-up analyses exploring the conditional effects of the predictor at values of the moderator revealed that the relationship between EE and RT was strongest in the older adults with shallower evidence accumulation build-up rates (Fig 5; CPP slope .0034, Coeff = -4.23, *SE*=1.17, *t*=- 3.61, *p*=.0009, 95% CI = [-6.60, -1.86]; CPP slope .0724, Coeff=-2.02, *SE*=.80, *t*=-2.52, *p*=.02, 95% CI = [-3.65 -.39], CPP slope .1405, Coeff =.16, *SE*=.98, *t*=.16, *p*=.87, 95% CI [-1.84, 2.16], Fig. 4). A post-hoc power analyses for the moderation model highlighted that power in excess of 99% was achieved (effect size Cohen’s *F*^2^=1.13, G*Power 3.1)

**Figure 4.**
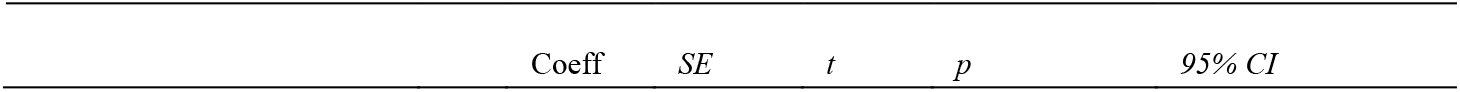
Moderation model demonstrating the relationship between EE and RT as moderated by CPP build-up rate. *Note* all analyses were conducted using continuous variables but are visualised here with three bins of equal size for CPP build-up rate.

**Figure 5.**
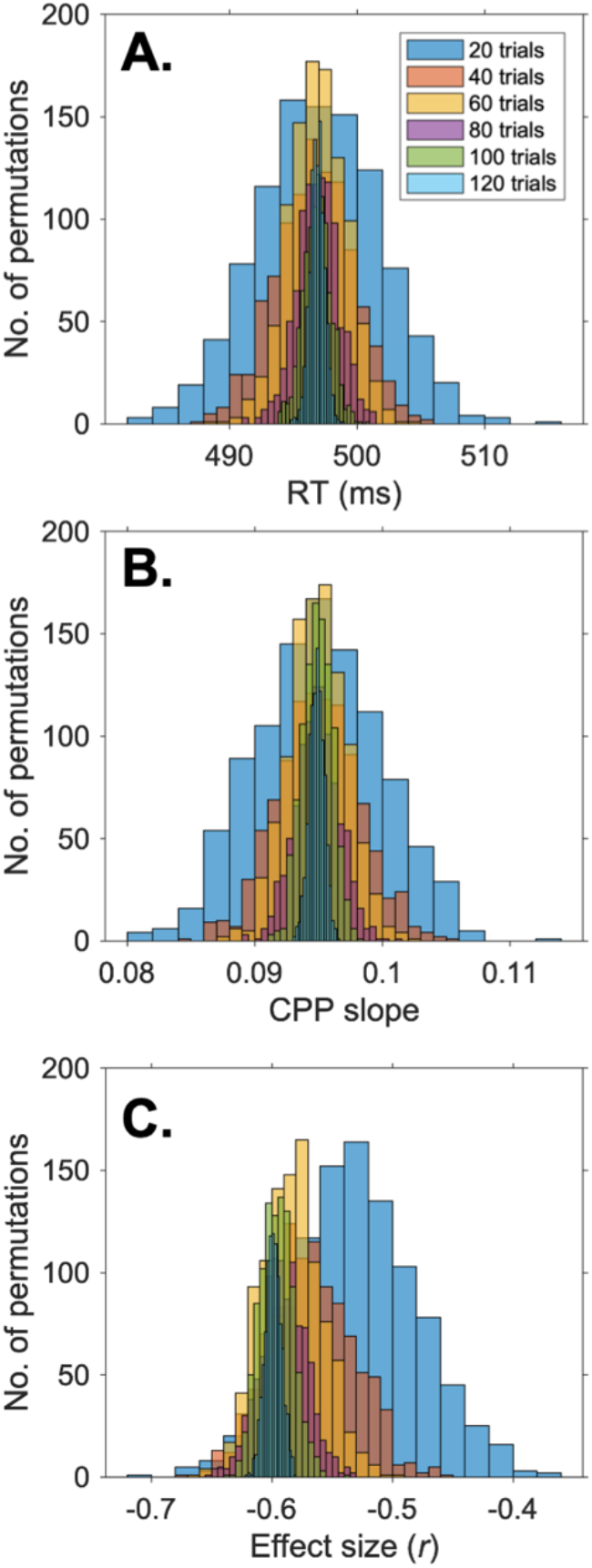
A-C. Reliable Estimates of the Relationship between RT and Evidence Accumulation Build-Up Rate Can be Obtained With Reduced Trial Numbers. For each participant we randomly selected 1000 estimates of RT (A) and CPP build-up rate (slope; B) for each of the six trial bin sizes (see legend). Reducing the number of trials reduced the signal to noise ratio and increased the likelihood that estimates of both RT and CPP build-up rate deviated from the *true* mean estimates (A, B). Critically, strong effect sizes (>.55) for the relationship between CPP build-up rate and RT were observed with as few as 40 trials (C) suggesting that this neurophysiological marker of sensory evidence accumulation may be developed as a translatable assessment of brain health for older adults.

**Table 3.**
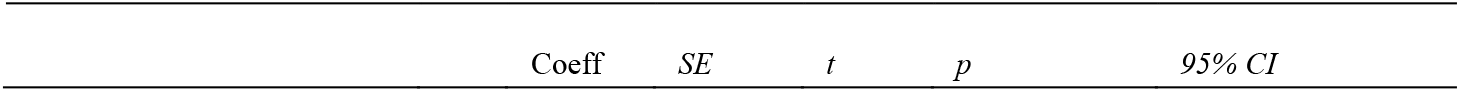
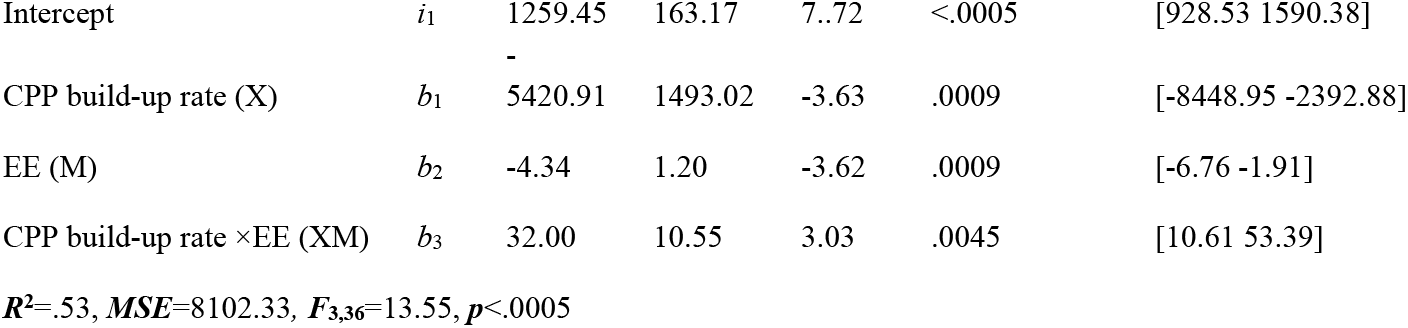
Results from a Regression Analysis Examining the Moderation of the Relationship between RT and Exposure to Environmental Enrichment in Older Adults by Neural Metrics of Evidence Accumulation Rate

These results suggest that the phenomenon of cognitive reserve, whereby high EE individuals are less reliant on typical markers of brain health to facilitate the preservation of cognitive function, is not just applicable to structural markers of brain health, but can be observed for neurophysiological markers. This provides a platform for future work to harness the millisecond temporal resolution of M/EEG to explore the neurophysiological basis of how this ‘reserve’ is facilitated. As such, these findings further suggest that the CPP build-up rate captures meaningful information relating to the neurophysiological health of the ageing brain.

### Feasibility of EEG markers of evidence accumulation build-up rate as a scalable proxy for neurocognitive health

Our findings provide evidence that the CPP build-up rate is mechanistically linked to an extensively validated marker of neurocognitive health – response speed – in older adults. This invites the possibility that this neural marker may be used by large-scale studies as an objective, cost-effective neurophysiological marker of ageing brain health. Both our results presented here, and a large body of previous research e.g. (Newman et al., 2017; McGovern et al., 2018; Steinemann et al., 2018; Brosnan et al., 2020), has measured the CPP using a single electrode (most typically ‘electrode *Pz’*). This affords clear benefits for reliably assessing this signal using low-density electrode arrays with either in-lab or portable EEG systems. Determining the minimum number of trials that permits a reliable measurement of CPP parameters, such as the CPP build-up rate, is therefore crucial for facilitating eventual clinical translation.

To determine this, we performed an analysis with a subset of participants who completed at least 8 task blocks, all of whom had a minimum of 129 valid response-locked ERP trials (see methods). For each participant, we derived 1000 estimates of RT and CPP build-up rate for each of the six trial sizes (Fig. 5.A, B) and tested whether the likelihood that the estimates of RT and CPP build-up rate were more likely to deviate from the *true* mean estimates with reduced trial numbers. Repeated measures ANOVAS using trial bin size as the factor and signal to noise ratio as the dependent measure (see methods) revealed a significant main effect of bin size for both RT (*F*_5,4995_=3820.14, *p*<.0005, partial *η*^2^ =79, Fig. 5A) and CPP build-up rate (*F*_5,4995_=298.13, *p*<.0005, partial *η*^2^ =.23, Fig. 5B). In both cases the data were best explained by a linear fit (RT: *F*_1,999_=10900.12, *p*<.0005, partial *η*^2^ =.92, CPP build-up rate: *F*_1,999_=859.60, *p*=<.0005, partial *η*^2^ =.46), indicating that increasing the number of trials significantly improved the signal to noise ratio.

To verify this pattern of results, we ran Kolmogorov-Smirnov tests on the mean estimates of CPP build-up rate and RT to assess whether the cumulative distribution function (CDF) increased with each reduction in trial number. These results demonstrated that when the number of trials was reduced by 20 trials, the width of the distribution (CDF) changed, as can be observed in Fig 5. A, B; Extended Data Fig. 5-1. This pattern of results demonstrates the expected effect that by reducing the number of trials, we increase the likelihood that estimates of both RT and CPP build-up rate deviate from the *true* mean estimates. Our critical question here, however, is at what level of SNR do we obtain reliable and behaviourally meaningful estimates of the relationship between RT and evidence accumulation build-up rate.

In the results reported in the main body of the text, we demonstrated a large effect size for the relationship between CPP build-up rate and RT (Pearson’s *r*)= -.60. Cohen’s (1988) cut-off for a large effect size is .5. As such, we defined the minimum number of trials at which a reliable CPP estimate can be derived as the number at which we can observe a strong effect size (i.e., an effect size greater than .55) for the relationship between RT and CPP build-up rate. To investigate this, we calculated the direct relationship, using Pearson’s correlation, between CPP build-up rate and RT for each of the 1000 permutations for each of the 6 bin sizes (20 up until 120 trials; Fig 5C). We then ran a Bayesian one sample *t*-test to test whether the estimates of effect size (*r*) for each bin size were significantly larger than -.55.

We found infinite support for the alternative hypothesis that the effect sizes for the relationship between RT and CPP build-up rate with 120, 100, 80, 60, and 40 trials were larger than .5 (all BF_10_>2.314ξ10+63; see descriptive statistics Table 4). However, this was not the case for the estimates derived using 20 trials. Here, Bayes factor analyses revealed strong support for the null hypothesis (BF_10_=.002), i.e., that the estimates of effect size were not greater than -.55 (Fig 5C, Table 4, Extended Data Fig. 5-2). As such, these results indicate that 40 response-locked trials are the minimum number of trials that will allow for a reliable estimation of the CPP build-up rate / RT relationship. With current paradigm timings (allowing for both variability in behavioural performance and quality of the EEG data), 40 trials could be obtained in less than 8 minutes (Extended Data Fig. 5-2), highlighting the potential for isolating reliable EEG metrics of evidence accumulation over relatively short time scales.

**Table 4.** Effect sizes for the relationship between RT and CPP build-up rates for 6 different trial sizes. Only at 20 trials did Bayes factor analyses reveal strong support for the null hypothesis that estimates of effect size were not greater than .5.

| $r$ CPP build-up rate-CPP | Mean | SD | $BF_{10}$ |
| --- | --- | --- | --- |
| 20 trials | -0.530 | 0.051 | .002 |
| 40 trials | -0.567 | 0.035 | $1.014 \times 10^{+45}$ |
| 60 trials | -0.584 | 0.025 | $2.829 \times 10^{+221}$ |
| 80 trials | -0.591 | 0.019 | ∞ |
| 100 trials | -0.596 | 0.014 | ∞ |
| 120 trials | -0.599 | 0.007 | ∞ |

### The Relationship between hDDM parameters and EEG markers of decision making

#### Non-decision time (t)

None of the EEG signals significantly improved the model fit for the non-decision time (*t*) derived using the hDDM (Extended Data Fig. 6-1). Thus, no evidence was found for a clear relationship between the neural metrics of decision making isolated using EEG and the *t* parameter of the hDDM.

**Figure 6.**
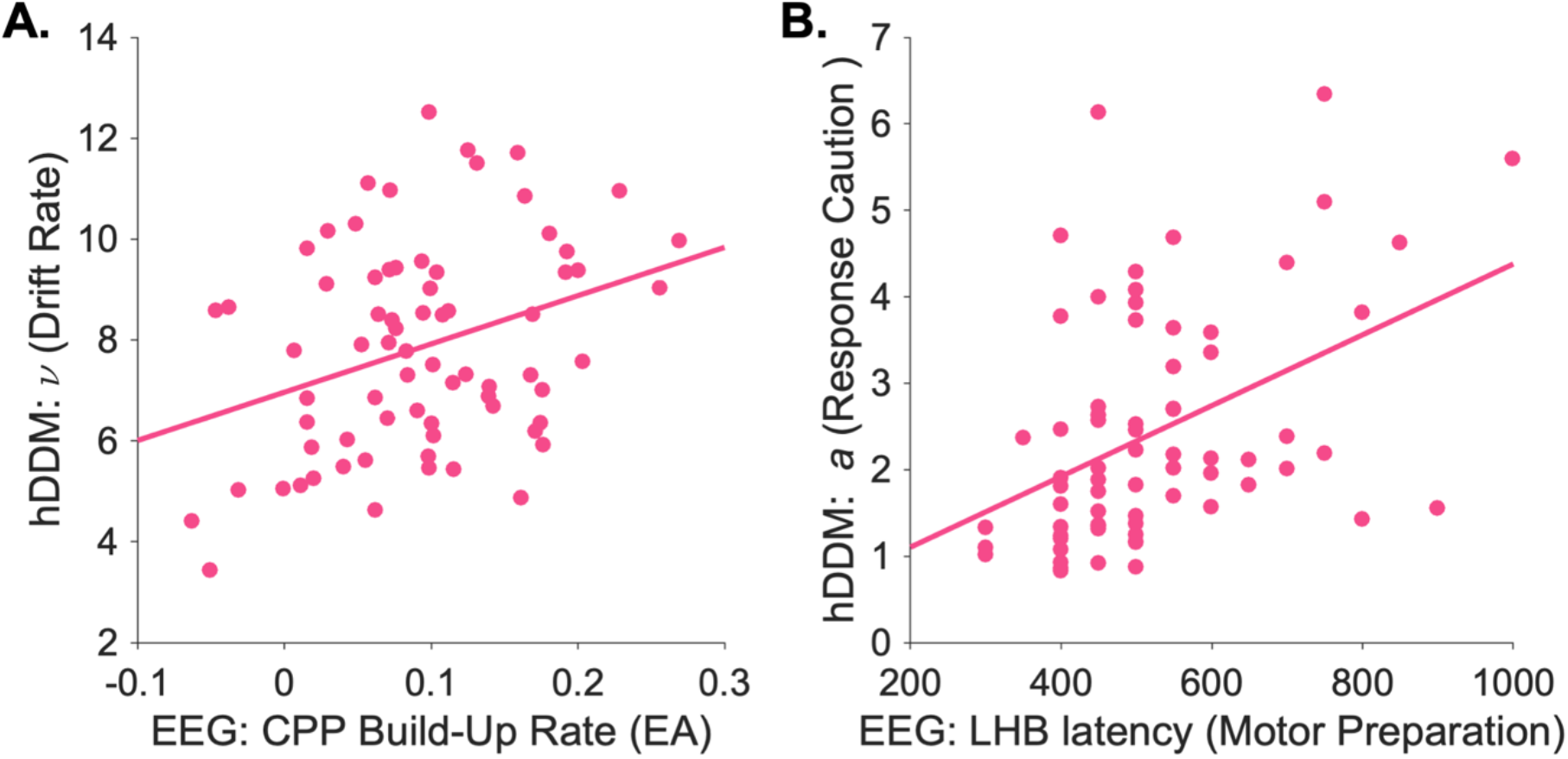
**A** Scatterplot of the association between CPP build-up rate, measured using EEG and drift rate (ν) isolated using a hDDM. **B.** Scatterplot of the association between LHB latency, measured using EEG and response caution (*a*) isolated using a hDDM.

#### Drift rate (ν)

In contrast, the model fit for drift rate (ν) was significantly improved when both age (Adjusted *R*^2^=.06, *F* Change = 5.74, *p*=.01) and the build-up rate of sensory evidence accumulation (CPP build-up rate; Adjusted *R*^2^=.08, *F* Change = 4.76, *p*=.03) were added to the model (Extended Data Fig. 6-2 for the full results from the hierarchical modelling procedure). To obtain accurate parameter estimates for the association between drift rate (ν) and CPP build-up rate, not influenced by other noninformative signals, Age and CPP build- up rate were entered into a separate linear regression model of drift rate (ν). This model explained 12% of the variance in drift rate (*F*_2,68_=5.78, *p*<.005; Parameter Estimates for CPP build-up rate: Standardised β = .28, *t*=2.34, *p*=.022; Age: Standardised β =-0.2, *t*=-1.57, *p*=.12, Extended Data Fig. 6-3, Fig. 6A).

#### Response caution (a)

The model fit for response caution (*a*) was significantly improved by the inclusion of both CPP amplitude (Adjusted *R*^2^= .14, *F* Change = 6.74, *p*=.01) and LHB Latency (Adjusted *R*^2^ = .22, *F* Change = 7.39, *p*=.0008; Extended Data Fig. 6-4). To obtain accurate parameter estimates for the association between response caution (*a*), CPP amplitude, and LHB latency, not influenced by other noninformative signals, Age, CPP amplitude, and LHB latency were entered into a separate linear regression model of response caution (*a*), Extended Data Fig. 6-5. This model explained 21.7% of the variance of response caution and only age and LHB latency (Fig. 6B) accounted for independent variation in response caution (*a*; *F*_3,67_=7.46, *p*<.001). Parameter Estimates: Age: Standardised β = .24, *t*=2.09, *p*=.04; CPP amplitude Standardised β = .20, *t*=1.83, *p*=.07; LHB Latency Standardised β = .38, *t*=3.35, *p*=.001, Fig. 6B).

#### The influence of age on the hDDM parameters

Older adults had lower drift rates (*ν* parameter; t_70_=-2.35, *p*=.02, 95% CI [-2.06 -.18]; older adults *M* =7.38, *SD*=2.05; younger adults *M*=8.49, *SD*=1.99), higher response caution (*a* parameter; *t*_70_=2.56, *p*=.01, 95% CI [.17 1.40]; older adults *M*=2.77, *SD*=1.54; younger adults *M*=1.98, *SD*=.88), and did not differ in non-decision time (*t* parameter; *t*_70_=.87, *p*=.39; older adults *M*=.11, *SD*=.06; younger adults *M*=.10, *SD*=.03).

## DISCUSSION

Here, we provide direct support for the hypothesis that build-up rates of sensory evidence accumulation are a critical neurophysiological mechanism underpinning the preservation of response speed in older adults. First, sensory evidence accumulation was not only directly related to response speed in older adults but also indirectly impacted performance by modulating subsequent neurophysiological processes, namely the decision criterion and the timing of the motor response. Second, consistent with the concept of neurocognitive reserve, a lifetime of EE offset age-related deficits in response speed. Critically, CPP slope moderated this association, such that the mitigating influence of EE on age-related declines in response times was most pronounced for individuals with relatively less efficient evidence accumulation (shallower build-up rates of CPP). This suggests that evidence accumulation build-up rates may offer rich information about which older individuals may benefit most from engaging with enriched environments.

In the current study, we saw concordance between the EEG and hDDM approaches for estimating sensory evidence accumulation such that higher drift rates (*v*) were associated with steeper CPP build-up rates. In addition, older adults presented with both lower drift rates and shallower EEG evidence accumulation build-up rates compared with their younger peers, further indicating that the rate of decision formation is disrupted with age. These results accord with previous work showing that the CPP closely conforms to the dynamics predicted by sequential sampling models (e.g. Twomey et al (2015) *European Journal of Neuroscience*; Kelly, Corbett & O’Connell (2021) *Nature Human Behaviour*), including in older populations (McGovern et al (2018) *Nature Human Behaviour*).

Deriving drift rate from the hDDM holds distinct practical clinical advantages over isolating the CPP using EEG, given that this parameter can be isolated from behavioural data alone. An interesting question for future longitudinal work will be to assess the utility of drift rate (ν) as an index of cognitive decline. In particular, it will be important to address whether this parameter provides more sensitivity as a clinical indicator of cognitive difficulties than behavioural assays of response speed alone, or markers of evidence accumulation rate isolated with EEG. In accordance with our findings, previous work has shown age-related differences in drift rate during random-dot motion task (McGovern et al., 2018). Yet McGovern and colleagues observed no differences in drift rate on a contrast change task in the same cohort of older adults. As such, an important avenue for future work will be to determine the extent to which drift rate (ν) is a *general* index of response speed in ageing as opposed to a metric that is specific to particular cognitive scenarios and tasks (see (Dully et al., 2018) for a review).

A key insight from decision modelling work with older adults has been that slowed RTs may not relate purely to sluggish information processing but might actually reflect a strategic preference for greater caution reflected in higher decision bounds (e.g. (Ratcliff et al., 2004, 2006a, 2006b)). These previous modelling studies have interpreted age-related increases in response caution (*a*), derived using drift diffusion modelling, as a more conservative decision threshold (and greater strategic preference for caution). In line with previous findings older adults did show greater response caution (*a*) compared with younger adults on the hDDM. However, our combined EEG-modelling results here suggest that independent variation in *a* is captured by the timing of the motor response (LHB latency) and not a higher decision threshold (CPP amplitude). In addition, older adults showed a lower CPP amplitude suggestive of a lower decision threshold. As such, our results show at least under the current task demands, age-related changes in response caution arise not from a rise in the decision threshold, but rather from slower preparation of the motor response. Although neural metrics of both the decision bound and the timing of motor preparation accounted for independent variation in response speeds, these relationships were contingent on the build-up rate of the CPP, such that slower build up rates of sensory evidence corresponded to lower decision bounds and longer response preparation speeds. Together these findings further indicate that response speed deficits obtained on an easy detection task in older adults result from a core deficit in the formation of perceptual decisions, as opposed to a more cautious approach to the decision-making process.

Although the collection and analysis of EEG data can be relatively time consuming, the data confer several distinct advantages that cannot be gleaned from behavioural or modelling alone. Through measuring brain activity with EEG, it is possible to distinguish the evidence accumulation process from other neurophysiological processes such as earlier sensory selection and later motor preparation processes, thereby offering unique insight into the neurobiology of age-related response speed slowing. In clinical cohorts, the same behavioural deficit (i.e., slowed response speed) can arise from a multitude of disrupted neural processes. By using EEG one can disentangle various processes in the brain and begin to identify the precise locus of dysfunction in clinical disorders such as stroke, Parkinson’s Disease and multiple sclerosis. This information about the brain systems and markers can then be used to assess deficits and develop novel interventions. Moreover, this is the first study to our knowledge to demonstrate the phenomenon of cognitive reserve outside of the imaging modalities, thereby providing a means to investigate the precise mechanisms by which EE alters brain function (as opposed to structure) in future neurophysiological research.

Future work may shed further light on the relationship between the formation of perceptual decisions and the motor response by incorporating measurements with equivalent millisecond precision. For example, subthreshold changes in the effector might be measured using continuous response measures such as voltage changes in hand-held force-sensing resistors (McBride et al., 2018) or changes in muscle activation with electromyography (e.g. Steinemann et al., 2018).

The results presented here are in keeping with the concept of cognitive reserve as defined by a recent consensus paper (Stern et al., 2020), but see (Cabeza et al., 2018, 2019; Stern et al., 2019), whereby the proxy of reserve (here EE captured by the CRIq) exerts a moderating influence on the relationship between markers of brain health and cognitive function (Stern et al., 2020). Our findings show that when evidence accumulation build-up rates are relatively shallower, individuals with relatively higher EE can nonetheless maintain faster response speeds than those with lower EE. One of the predominant principles of cognitive reserve is that high EE individuals are less reliant on established markers of brain health for facilitating behaviour. Our findings accord with a large body of work which has demonstrated the phenomenon of cognitive reserve with structural (e.g. grey matter atrophy in healthy individuals (Chan et al., 2018)), and neuropathological (e.g., amyloid plaques and tangles in Alzheimer’s patients (Xu et al., 2019)) markers of compromised brain health. Here, we show that cognitive reserve can also be observed using neurophysiological markers of ageing.

Although our results provide evidence that cognitive reserve can be indexed using EEG, the mechanisms supporting cognitive resilience in high EE individuals is an avenue for future work (Cabeza et al., 2018, 2019; Stern et al., 2019, 2020). An important direction will be to capitalise on the current findings and harness neurophysiological techniques to understand the neurobiological substrates of cognitive reserve.

Fronto-parietal regions of the brain may contribute to preserved RT in higher EE individuals, even when they experience compromised markers of brain health. Converging evidence from structural MRI, resting state MRI, transcranial electrical stimulation, and post mortem histology in healthy older adults, patients with mild cognitive impairment, and individuals with Alzheimer’s disease points to a critical role for the frontal lobes, particularly the prefrontal cortex (PFC), in supporting resilience (Valenzuela et al., 2012; Brosnan et al., 2017, 2018a, 2022; Franzmeier et al., 2017a, 2017b; Shalev et al., 2020). The PFC (specifically the dorsolateral PFC) is the largest functional component of a domain-general ‘multiple demands’ system of fronto-parietal, and insular brain areas (Duncan, 2001, 2013) which is activated during a wide range of cognitive operations (Fedorenko et al., 2013). This system exert a ‘top-down’ modulatory influence over many brain areas and cognitive processes (Cristescu et al., 2006; Summerfield et al., 2006; Voytek et al., 2010; Nelissen et al., 2013; Brosnan et al., 2018a). It is possible that the multiple cognitive demands necessitated by enriched environments such as educational settings, complex occupational environments, social and leisure activities continuously require recruitment of this network which, over time, may benefit cognition. Accordingly, a number of studies have shown that connectivity within the fronto-parietal networks (FPN) accounts for substantial inter-individual variability in neurocognitive resilience in older adults (e.g. (Franzmeier et al., 2017a, 2017b; Veldsman et al., 2020; Brosnan et al., 2021)).

In younger adults, we have recently shown that neural metrics of evidence accumulation rate vary according to individual differences in connectivity within the dorsal FPN (white matter macrostructural organisation of the superior longitudinal fasciculus (SLF), and resting state functional connectivity within the dorsal FPN (Brosnan et al., 2020)). The results of current study provide an unprecedented framework for exploring whether higher EE individuals should show differences in structure and/or function in fronto-parietal regions to allow them maintain preserved response speeds despite suboptimal evidence accumulation capacities in future work. Using EEG in combination with techniques that allow examination of the FPN with more precise spatial specificity (diffusion MRI, functional MRI, and MEG) will be particularly useful for asking how higher EE individuals might preserve fast responses despite compromised evidence accumulation build-up rates. This would help disentangle the interacting influences of EE, fronto-parietal regions, and distinct stages of information processing and perceptual decision formation to cognition.

A question of pressing societal relevance is to identify enriched cognitive, social, and leisure environments with which older adults could engage in later in life to optimise resilience to cognitive decline. Several lines of evidence suggest the association between EE and cognition is causal. For example, monozygotic twin pairs exposed to greater levels of enrichment throughout life show relatively faster response speed in later years (Lee et al., 2014). Similarly, a three-month intervention learning new skills in healthy older adults improves response speed (Park et al., 2014). Finally, emerging results from a large (*N*=2832) multicentre longitudinal clinical trial using computerised speed-based cognitive training in older adults shows that training this may causally improve neurocognitive health in older adults (Ball et al., 2002, Rebok et al., 2014, Wolinsky et al., 2009, Wolinsky et al., 2009, Edwards et al., 2017). In our data, the association between EE and behaviour was driven exclusively by the leisure, and not education and occupation subscales, of the CRIq. These findings accord with growing evidence that leisure and social activities are critical for supporting brain health in ageing (e.g., Lee et al., 2014; Xu et al., 2018). This is of relevance to public health interventions given the accessible and potentially modifiable nature of leisure and social activities, particularly in later years of life. Further work using the CRIq in larger sample sizes to run item-level analyses would help understand precisely which type of leisure activities would optimally facilitate resilience. This in turn would pave the way towards developing scalable, affordable public health interventions to induce lasting, positive changes in ageing brain function and resilience cognitive decline. A limitation of the current design is that we did not measure socioeconomic status (SES), which is known to independently contribute to brain health and neurocognitive resilience (e.g., Jones et al., 2011). As such, it is not possible for us to disentangle the extent to which the association between EE with our neurocognitive markers is mediated by socioeconomic factors. It is important for future studies, using larger cohorts, to investigate this and identify facets of brain health that are modifiable, irrespective of SES, to increase resilience to cognitive decline.

Recently, there has been a focus on the motivational processes that govern how resources are allocated to effortful tasks (Chong, 2018; Westbrook and Frank, 2018; McGuigan et al., 2019; Westbrook et al., 2020). This idea is particularly relevant to our study given that older adults have been shown to outperform their younger counterparts on cognitively effortful tasks (Mather and Carstensen, 2005; Ennis et al., 2013). For example, older adults demonstrate a positivity bias resulting from enhanced cognitive control over positive emotions (Mather and Carstensen, 2005). In our study, the CRI data suggest that high EE individuals tended to engage in activities associated with higher levels of motivation relative to their low EE peers (e.g., social activities, attending conferences, exhibitions, and concerts, and using modern technology). Future work with larger cohorts could directly test whether motivation in older adults, and especially those with high EE, can overcome evidence accumulation deficits to facilitate fast responses.

Finally, identifying the precise stage of information processing driving slowed response speed with aging might hold valuable prognostic information and could provide a sensitive addition to future large-scale epidemiological and translational studies. In addition, our framework outlines a means to investigate the mechanisms by which high EE individuals may compensate for deficits in evidence accumulation to maintain fast responding. Future work should expand our targeted and comprehensive EEG analysis to explore the role of motivation and cognitive control in this regard. We present further evidence here that we can obtain reliable (large effect sizes) and meaningful (strongly predictive of response speed) measurements of the CPP build-up rate with as few as forty trials. Taken together, our work suggests that measuring the CPP via low density and potentially portable EEG, might have significant value for exploring the mechanisms by which EE positively benefits brain function.

These findings suggest that neural metrics of evidence accumulation build-up rate index an important facet of neurocognitive vulnerability in the ageing brain. Moreover, they suggest that CPP build-up rate holds promise as an EEG marker indexing a critical facet of neurophysiological vulnerability of the ageing brain that could be incorporated into large scale epidemiological studies. It will be important for future work to replicate these effects and interrogate mechanisms supporting the maintenance of fast response speed in higher EE individuals.

## EXTENDED DATA

**Extended Data Figure 2-1:**
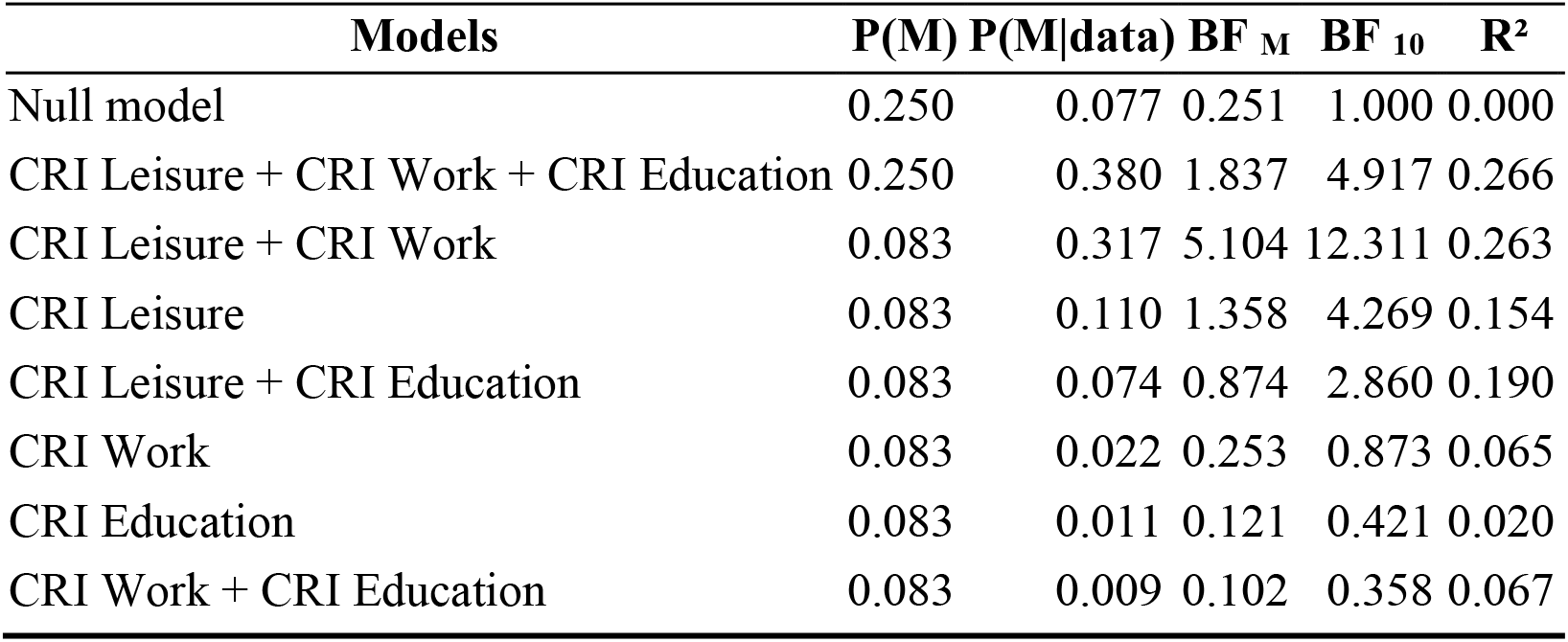
Bayesian Linear Regression Model the relationship between RT and Cognitive Reserve.

**Extended Data Figure 2-2.**
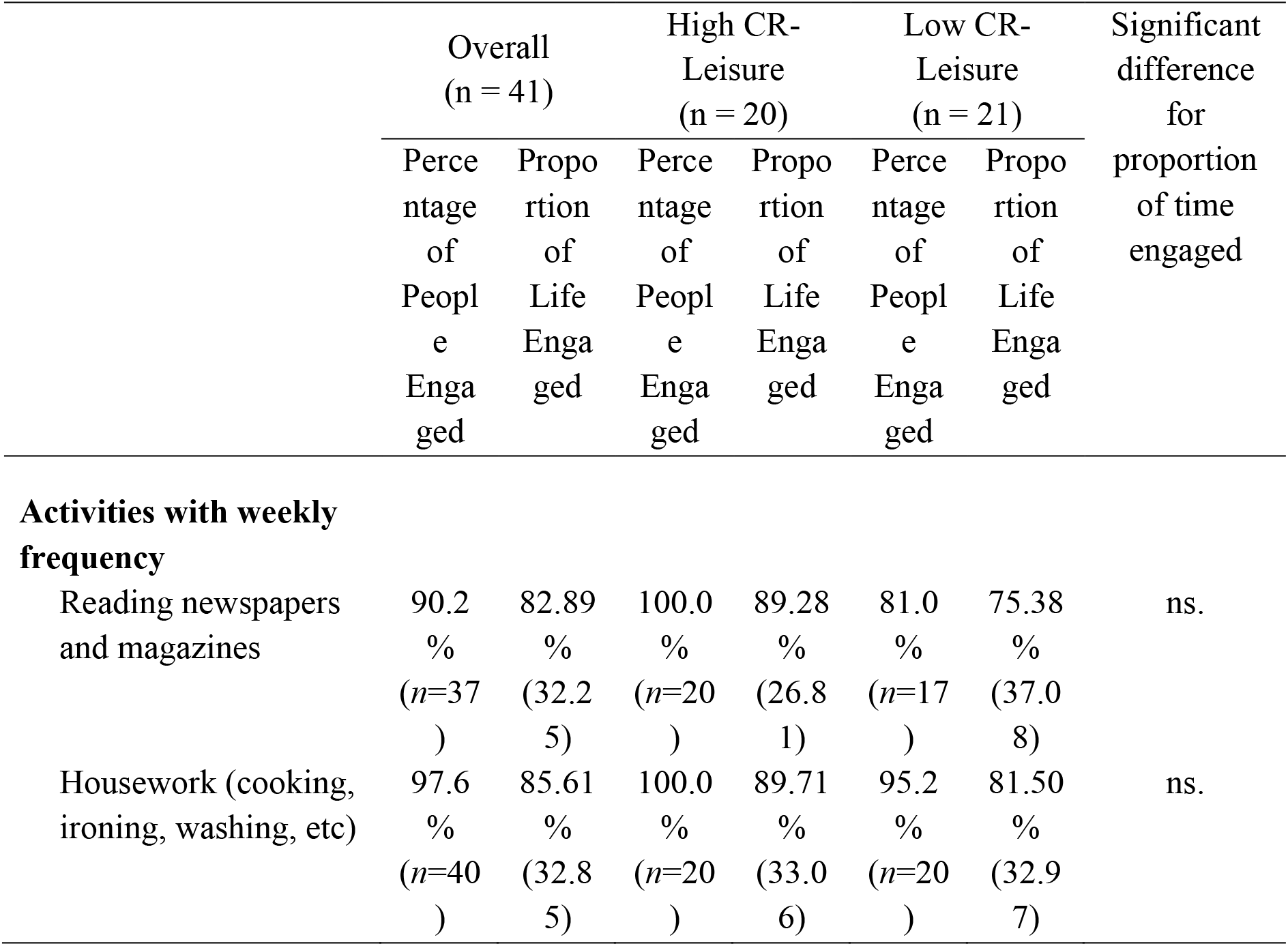
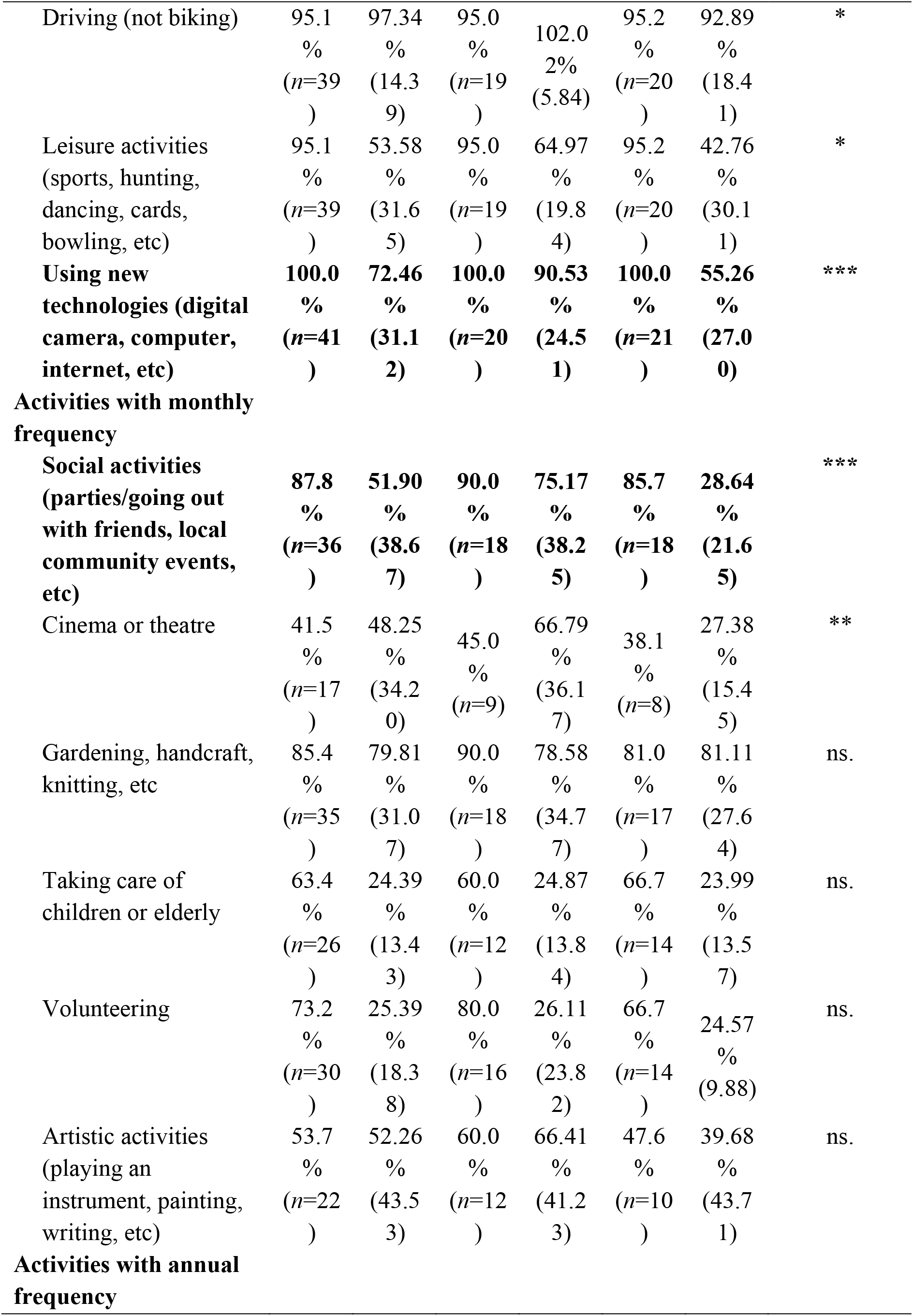
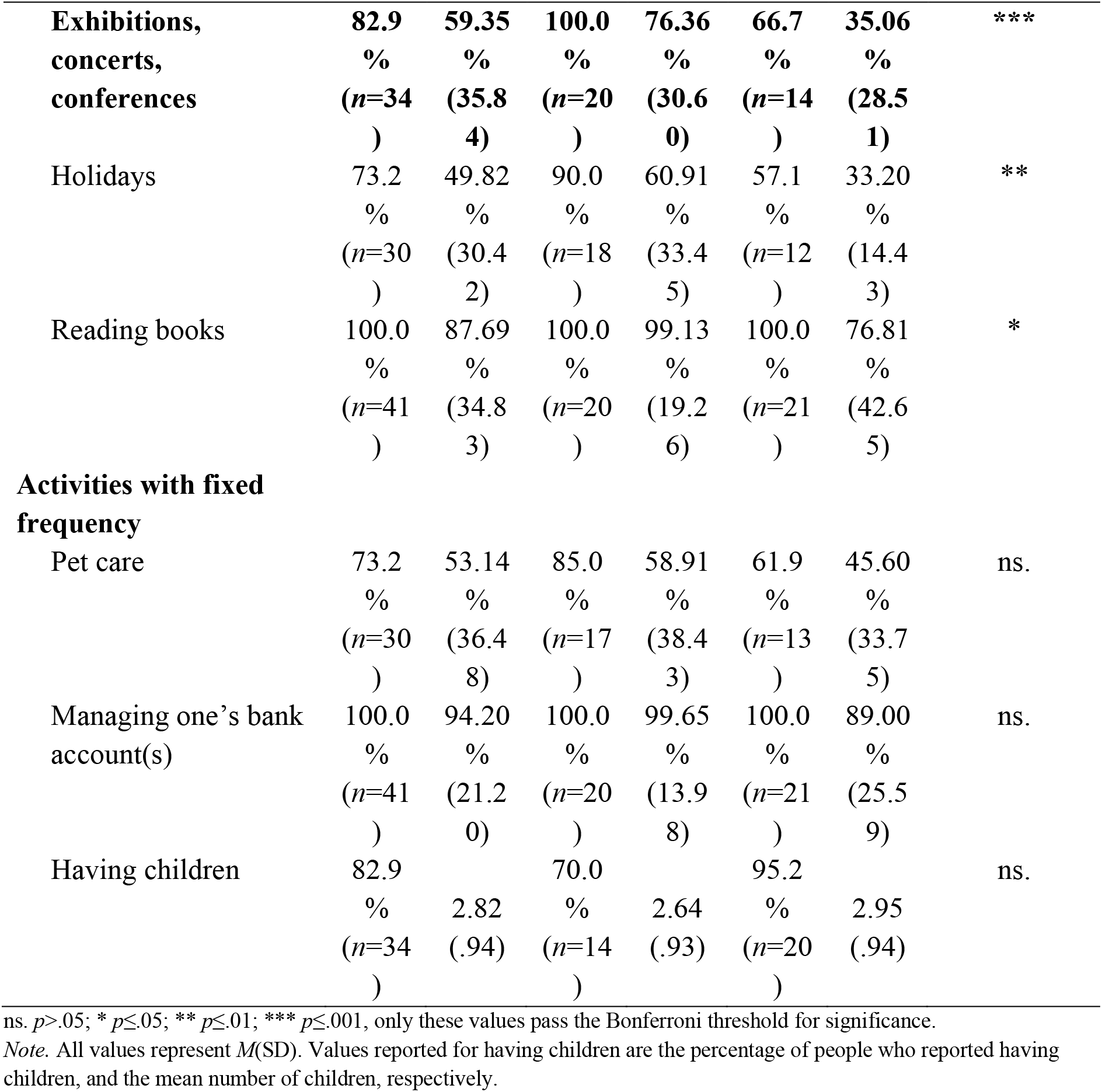
Frequency and duration of leisure activity engagement for those with high and low CRI Leisure (devised by median split).

**Extended Data Figure 2-3.**
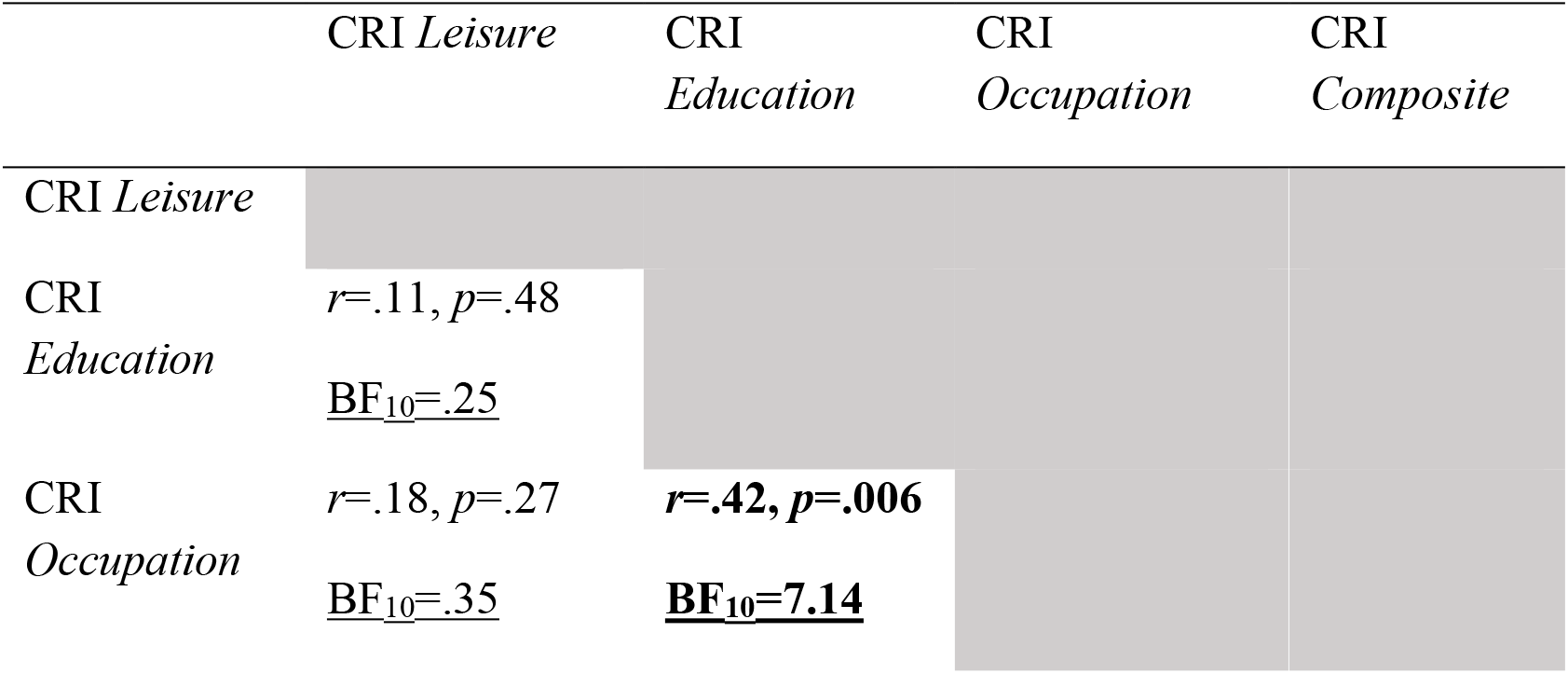
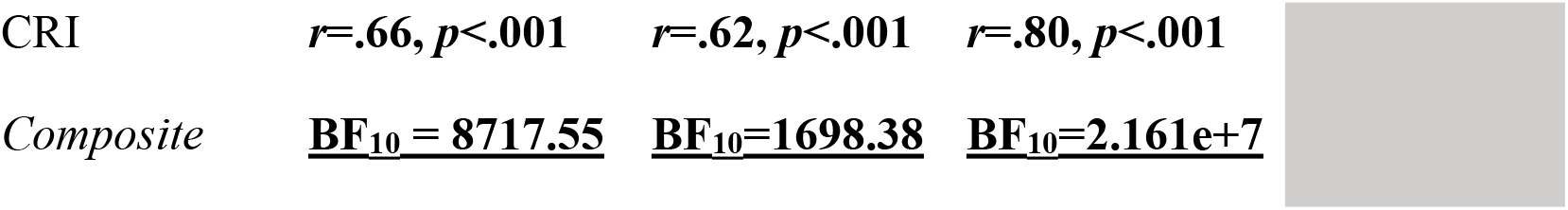
The effects of CRIq Leisure on RT are statistically independent to the association between RT and CRI Education and CRI Occupation.

To explore the association between the CRIq Leisure subscale with Education and Occupation, we ran correlational analyses between the four CRI subscales (Supplementary Table 8). First, the composite score was strongly associated with leisure, occupation, and education engagements (all *r*>.65), *p*<.001). A robust association was also observed between the CRI subscales of education and occupation (*r*>.4, see Table below). Critically, however, we did not find evidence to suggest an association existed between the CRI *Leisure* subscale with CRI Education (*r*=.11, *p*=.48) or CRI *Occupation* (*r*=.18, *p*=.27). In fact, follow up Bayesian analyses suggested moderate-anecdotal evidence in support of the null hypothesis, i.e., that no relationship existed between the Leisure subscale with Education (BF10=.25) or Occupation (BF10=.35). As such, these results suggest that our measure of cognitive reserve driving the associations with RT (i.e., the CRI leisure subscale) is statistically independent of the education and occupation subscales.

To address the concern that our observed associations between RT and CRI Leisure could be attributed to the effects of education or occupation, we re-ran the regression models while controlling for the effects of education and occupation.

First, we re-ran the final regression model, in which response speed was modelled as a function of the CRI Leisure subscale, this time adding both the CRI Education and CRI Occupation subscales as nuisance covariates in step 1. With this approach, we observed that modelling RT as a function of CRI Education and CRI Occupation did not account for significant variance in RT (Adjusted *R*^2^=.017, *F*_(2,38)_=1.34, *p*=.27). Critically, when the CRI Leisure subscale was added to the next stage of the model, there was a significant improvement in model fit, over and above the null model (Adjusted *R*^2=^.203, *R*^2^ change = .197, *F* change = 9.89, *F*_(3,37)_=4.40, *p*=.01; <u>Standardized *β* CRI Leisure= -0.45, *t*=-3.15, *p*=.003; 95% CI [-4.97 -1.07]</u>; Standardized *β* CRI Education= 0.06, *t*=0.41,, *p*=.66; 95% CI [-2.71 4.08]; Standardized *β* CRI Occupation= .31, *t*=1.95, *p*=.06; 95% CI [-.07 3.91]).

Second, to verify our results using Bayesian statistics, we re-ran the Bayesian Linear Regression model, this time adding the CRI subscales of Education and Occupation to the null model, thereby exploring the direct association between the CRI Leisure and RT. This analysis indicated strong evidence in support of the hypothesis that CRI Leisure accounts for substantial variance in behavioural markers of response speed (BF_10_=14.38). As such, these results suggest our effects are driven by leisure activities, and not education or occupational engagements.

**Extended Data Figure 2-4.**
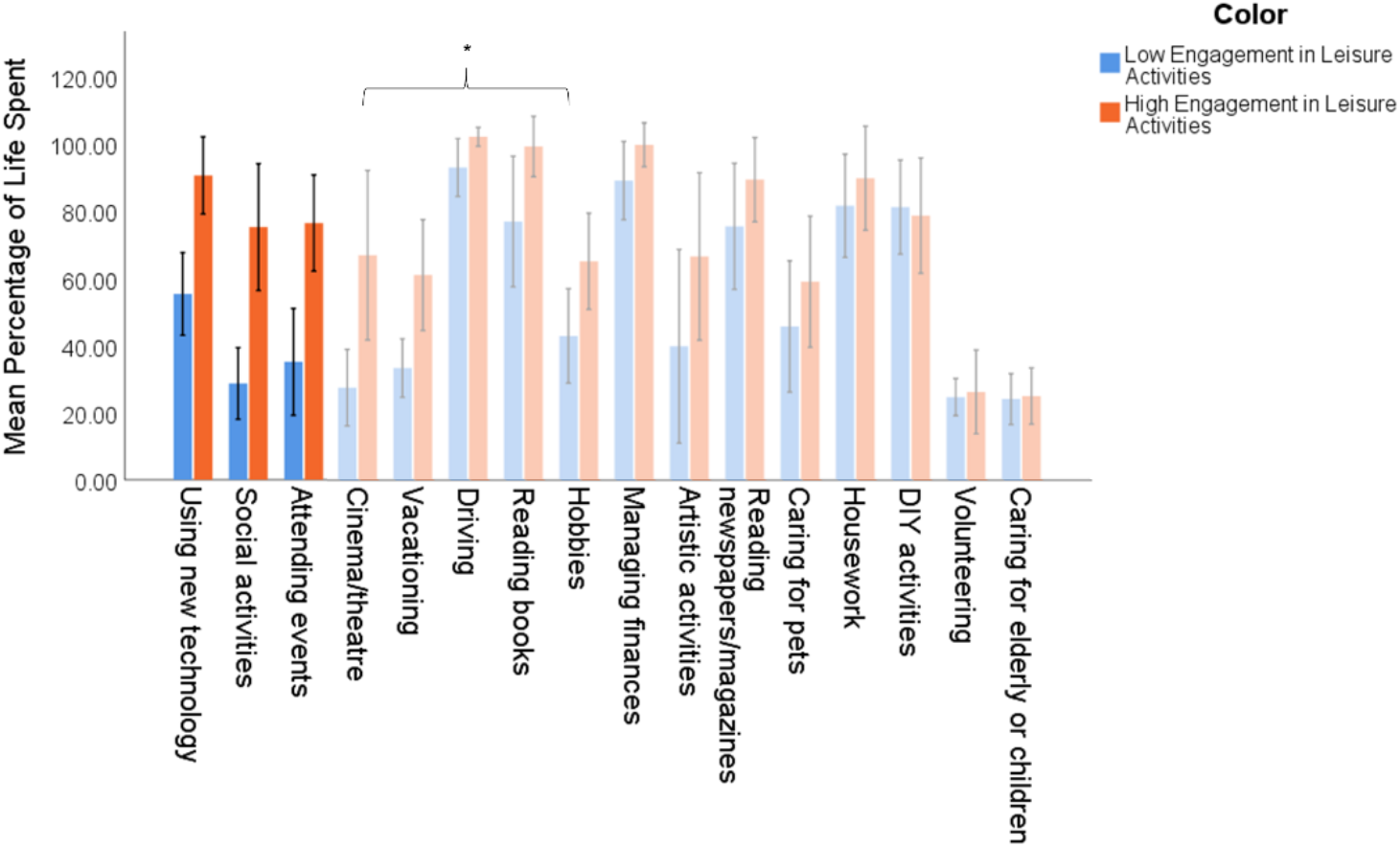
Differences between the activities of those with relatively higher versus lower levels of lifetime leisure engagement. Proportion of life spent engaging in particular activities varies between those with an overall higher or lower level of engagement in leisure activities. Significant group differences are presented in bold. * denotes comparisons where *p*<.05, but did not satisfy a Bonferroni adjusted α=.003.

To explore the leisure activities which may drive the apparent effect of EE on response speed, we compared the activities of those with higher versus lower levels of lifetime leisure engagement. To do so, we first devised two groups of older adults based on a median split of their engagement in leisure activities. Those with CRI *Leisure* subscores above the overall median score of 138.00 were considered *High Engagement* (*n*=20), while those with a subscore equal to or lower than 138.00 were deemed *Low Engagement* (*n*=21). We then examined each participant’s responses to individual activities on the CRIq.

Participants first indicated whether they participated in the activity *Often/Always*, or *Never/Rarely* over the course of their lifetime, and further specified for how many years they engaged *Often/Always*. For participants who engaged in an activity *Often/Always* for at least one year of life, we calculated separate values for their engagement in each activity, representing the percentage of life years spent engaging in each activity since 18 years of age using the following formula: ((Years of activity)/(Age – 18)) * 100. Note that rounding within the CRIq causes some individuals to exceed 100.00% of life spent participating in a given activity. For example, if an individual worked as a nurse for 3 years, this is rounded up to 5 years, as per the standardised questionnaire administration guidelines. Finally, we determined the percentage of individuals in each group who engaged in each activity *Often/Always*, and the mean percentage of life spent engaging in each activity for each group, the results of which are demonstrated in Extended Data Figure 2-2.

Subsequently, we investigated group differences through a series of *t* tests. Those with *High Engagement*, compared to those with *Low Engagement* in leisure activities, spent a significantly greater proportion of their lives using modern technology (*t*_39_=4.37, *p*<.001), engaging in social activities (*t*_26.88_=4.49, *p*<.001), and attending events such as conferences, exhibitions and concerts (*t*_32_=3.98, *p*<.001), vacationing (*t*_24.83_=3.11, *p*=.01), cinema or theatre attendance (*t*_11.09_=2.85, *p*=.01), driving (*t*_22.96_=2.11, *p*=.05), reading books (*t*_28.12_=2.18, *p*=.04), or engaging in hobbies such as sports and games (*t*_37_=2.31, *p*=.03). No other significant differences were found (all *p*>.05).

We additionally ran a series of Bayesian independent samples *t* tests to compare our ‘high leisure’ and ‘low leisure’ groups on each of the 16 leisure activities. These suggested that our high leisure group spent more of their lives attending events such as conferences, exhibitions, and concerts (BF_10_=10658.60), using modern technology (BF_10_=240.49), engaging in social activities (BF_10_=106.02), and vacationing (BF_10_=72.46). The BF_10_ of all other models was <3.00 indicating at most anecdotal evidence in support of the alternative hypothesis.

**Extended Data Figure 2-5.**
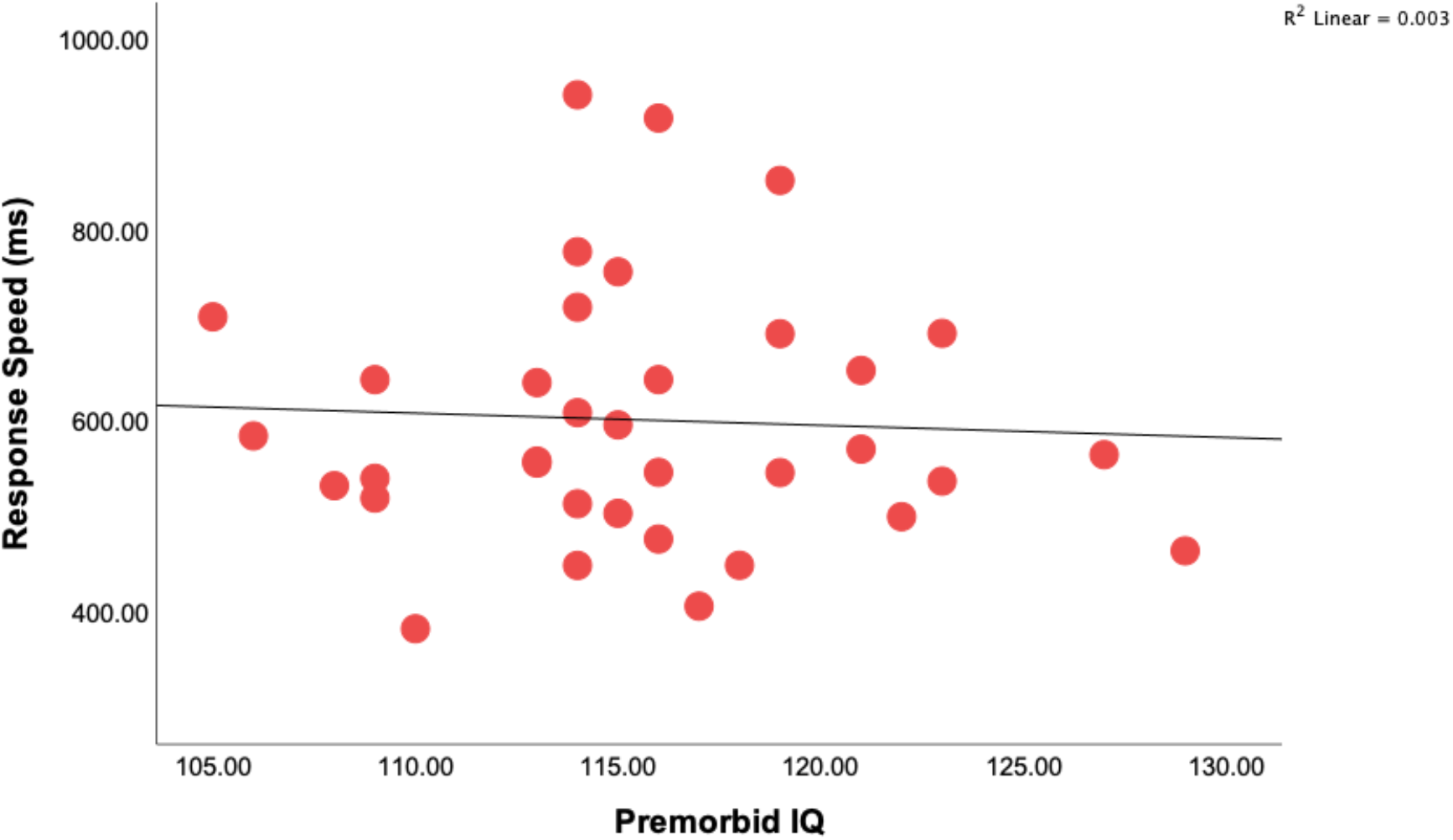
The impact of IQ on Response Speed. To investigate whether the observed relationship between EE and response speed could reflect individual differences in IQ, we estimated premorbid intelligence in a subset of participants (n=36, not shown). There was no direct association between IQ and response speed. Critically, the relationship between EE and response speed remained significant after covarying for IQ, indicating that the relationship between enrichment of cognition was not due to individual differences in intelligence. More specifically, a subset (*N*=36) of the older adults completed word reading tasks commonly used to estimate premorbid IQ based on the Wechsler Adult Intelligence Scale – Fourth Edition (WAIS-IV) (Wechsler, 1981). Of these individuals *n*=17 completed the Test of Premorbid Function (ToPF), while the other *n*=19 completed the National Adult Reading Test (NART) (Nelson, 1982), using updated norms. Outliers were defined in SPSS using the IQR, consistent with the main analyses, separately for both cohorts of older adults. One outlier was detected for the ToPF and was subsequently removed and imputed using the mean value from their group. The two cohorts differed on estimates of IQ score derived using the different word lists (*t*_34_=2.09, *p*=.04), with a subsequent Bayesian independent samples *t* test suggesting moderate evidence for a difference between the two groups, BF_10_ =3.26. This is likely attributable to established differences in the estimations produced by the measure. Nonetheless, we considered it useful to investigate using the data available to us, whether our effects could be attributed to a relationship between response speed and IQ. For this, we ran a hierarchical linear regression of RT, with IQ entered as the first step in the model. IQ did not account for a statistically significant proportion of the variance in RT, indicating no direct influence of premorbid intelligence on response speed (*F*_(1,34)_ =.07, *p*=.80, *R^2^* =-.03). Critically, when the CRI sub-scales were added to the second step in this hierarchical model, the relationship between EE and response speed remained significant (*R_adj_^2^* =.17, *R_changej_^2^* =.26, *p*=.02; *F*_(4,31)_=2.77, *p*=.04), demonstrating that the observed relationship between EE and response speed cannot be attributed to IQ. Note that due to changes in the protocol for a larger clinical (stroke) study under which these data were collected our measurement of IQ changed from the NART to TOPF midway through this clinical project. As such, the results here should be interpreted with caution given the two different IQ measurements used.

**Extended Data Table 2-1.** *Note* all values signify mean (M) and standard deviations (SD). EEG signals that differed significantly between the older and younger adults (according to both Bonferroni-corrected frequentist and Bayesian analyses) are highlighted in bold. *Note*, weak (anecdotal) evidence is provided for the group difference in N2c amplitude

| | Younger Adults<br>($N=31$) | Older Adults<br>($N=41$) |
| --- | --- | --- |
| N2c Amplitude ($\mu V/m^2$) | -10.03 (8.35) | -15.98 (11.36) |
| N2c Latency (ms) | 266.16 (55.61) | 285.78 (44.81) |
| **CPP Onset (ms)** | **250.84 (80.39)** | **345.17 (117.16)** |
| **CPP Build-up Rate** | **.12 (.07)** | **.07 (.07)** |
| CPP Amplitude ($\mu V/m^2$) | 24.32 (13.75) | 17.20 (11.16) |
| LHB Latency (ms) | 471.58 (98.18) | 549.02 (178.46) |
| LHB Slope | -.0059 (.0057) | -.0068 (.0067) |
| LHB Amplitude | -1.32 (1.26) | -2.04 (1.73) |

**Extended Data Table 2-2.** Hierarchical linear regression model statistics examining how each neurophysiological marker contributed to RT, over and above the contributions made by those processes that temporally preceded.

Hierarchical Modelling of RT
| Model |  | Sum of<br>Squares | df | Mean Square | F | Sig. |
| --- | --- | --- | --- | --- | --- | --- |
| **Age (centred)** |  |  |  |  |  |  |
| 1 | Regression | 440379.981 | 1 | 440379.981 | 40.705 | .000<sup>b</sup> |
|  | Residual | 746497.607 | 69 | 10818.806 |  |  |
|  | Total | 1186877.588 | 70 |  |  |  |
| **+ N2c Latency** |  |  |  |  |  |  |
| 2 | Regression | 457738.096 | 2 | 228869.048 | 21.344 | .000<sup>c</sup> |
|  | Residual | 729139.492 | 68 | 10722.640 |  |  |
|  | Total | 1186877.588 | 70 |  |  |  |
| **+ N2c Amplitude** |  |  |  |  |  |  |
| 3 | Regression | 474126.961 | 3 | 158042.320 | 14.856 | .000<sup>d</sup> |
|  | Residual | 712750.627 | 67 | 10638.069 |  |  |
|  | Total | 1186877.588 | 70 |  |  |  |
| **+ CPP Onset Latency** |  |  |  |  |  |  |
| 4 | Regression | 477414.083 | 4 | 119353.521 | 11.103 | .000<sup>e</sup> |
|  | Residual | 709463.505 | 66 | 10749.447 |  |  |
|  | Total | 1186877.588 | 70 |  |  |  |
| **+ CPP Build-Up Rate** |  |  |  |  |  |  |
| 5 | Regression | 675255.405 | 5 | 135051.081 | 17.158 | .000<sup>f</sup> |
|  | Residual | 511622.183 | 65 | 7871.111 |  |  |
|  | Total | 1186877.588 | 70 |  |  |  |
| **+ CPP Amplitude** |  |  |  |  |  |  |
| 6 | Regression | 757783.795 | 6 | 126297.299 | 18.837 | .000<sup>g</sup> |
|  | Residual | 429093.793 | 64 | 6704.591 |  |  |
|  | Total | 1186877.588 | 70 |  |  |  |
| **+ LHB Build-Up Rate** |  |  |  |  |  |  |
| 7 | Regression | 757824.042 | 7 | 108260.577 | 15.896 | .000<sup>h</sup> |
|  | Residual | 429053.546 | 63 | 6810.374 |  |  |
|  | Total | 1186877.588 | 70 |  |  |  |
| **+ LHB Peak Latency** |  |  |  |  |  |  |
| 8 | Regression | 816914.302 | 8 | 102114.288 | 17.113 | .000<sup>i</sup> |
|  | Residual | 369963.286 | 62 | 5967.150 |  |  |
|  | Total | 1186877.588 | 70 |  |  |  |
| **+ LHB Amplitude** |  |  |  |  |  |  |
| 9 | Regression | 828221.160 | 9 | 92024.573 | 15.651 | .000<sup>j</sup> |
|  | Residual | 358656.428 | 61 | 5879.614 |  |  |
|  | Total | 1186877.588 | 70 |  |  |  |

**Extended Data Figure 3-1:**
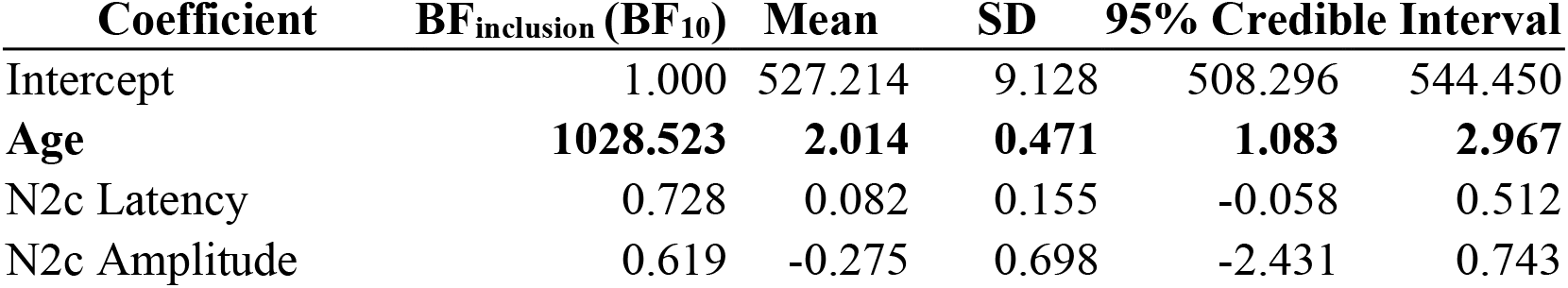
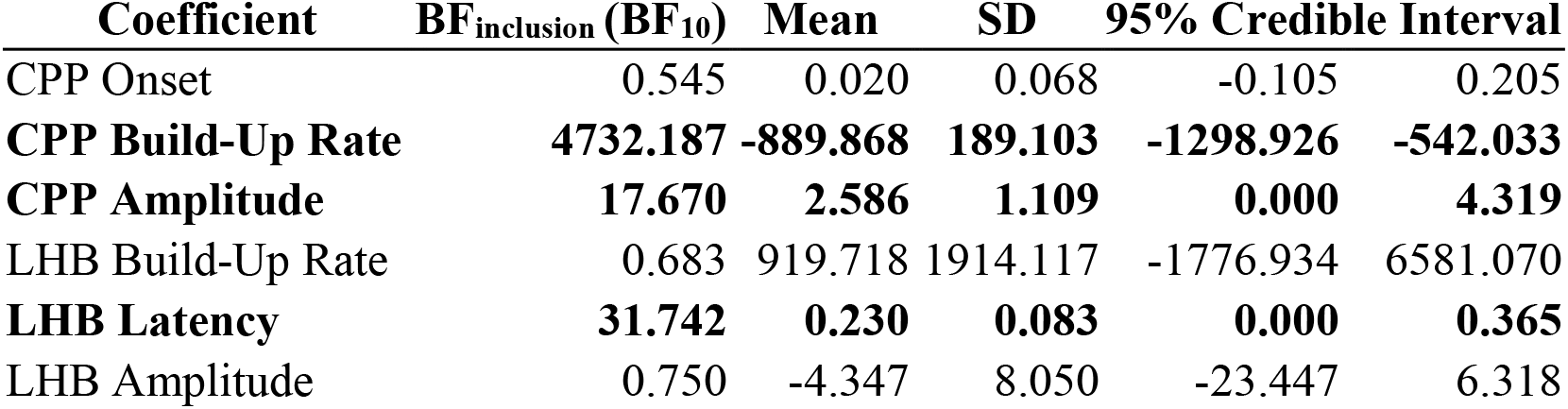
Bayesian linear regression model statistics examining how each neurophysiological marker contributed to RT. Note. BF_inclusion_ values above 1 indicate the strength of evidence in favour of the alternative hypothesis and are highlighted in bold.

**Extended Data Figure 3-2.**
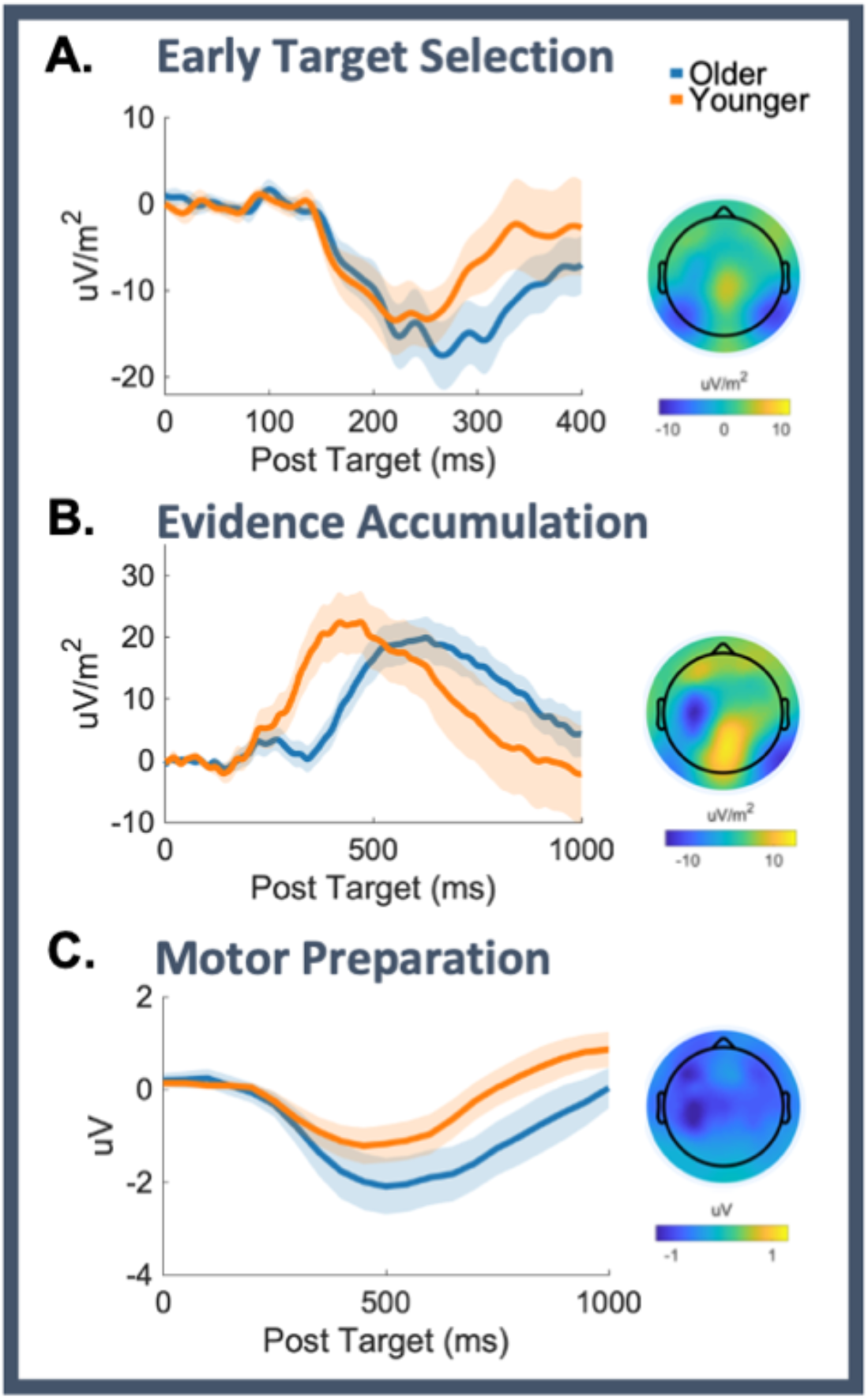
Temporal Dynamics of Evidence Accumulation Are Robust Age-Related Indicators. **A**. The stimulus-aligned N2c waveform (electrodes P7/P8) for older and younger adults. **B.** Stimulus-aligned CPP waveform (electrode Pz) for the two groups. **C.** Stimulus-aligned beta waveform (electrode C3) for the two groups. *Note*. The topoplots depict the spatial distribution of the EEG signal for both groups combined at 150- 400ms post-target for the N2c (**A**) -150ms to 50ms aligned to response for the CPP **(B)** and 400- 700ms post-target for LHB **(C).** We examined group-level differences in the eight electrophysiological markers using a series of one-way ANOVAs, *Bonferroni*-corrected for multiple comparisons (alpha .05/8 EEG components => alpha-corrected threshold = .006) and supplemented these with Bayesian analyses to indicate the strength of evidence in support of the null hypothesis. No statistically significant difference was observed between older and younger adults in the latency of early target selection signals (N2*c;* F_1,70_ =2.75, *p*=.10, BF_10_=.79, Supplementary Table 1 for plots and additional analyses see Supplementary Fig. 2), and although there was weak evidence to suggest that the amplitude of the N2c differed between groups (*F*_1,70_ =6.01, *p*=.02, partial *η*^2^=.08, BF_10_=3.05, Supplementary Table 1, Supplementary Fig. 1A), this did not survive correction for multiple comparisons. In line with recent reports(McGovern et al., 2018), the older adults differed from their younger counterparts in metrics of evidence accumulation (the CPP). More specifically, timing delays were observed for several parameters of the CPP in older individuals; they showed a later onset (<u>later CPP onset</u>; *F*_1,70_=14.8, *p*<.001, partial *η*^2^=.18, BF_10_=96.52, Supplementary Table 1, Supplementary Fig. 1B) and slower build-up rate (<u>shallower CPP build-up rate</u> (slope); *F*_1,70_ =8.03, *p*=.006, partial *η*^2^=.10, BF_10_=6.90, Supplementary Table 1, Supplementary Fig. 1B). No groups differences were observed for the amplitude at response (CPP amplitude; *F*_1,70_=5.88, *p*=.02, partial *η*^2^=.08, BF_10_=2.89, Supplementary Table 1). We did not observe differences in motor preparatory activity between the two groups (LHB build-up rate (slope) *F*_1,70_=.31, *p*=.58, BF_10_=.28; stimulus-aligned peak LHB latency *F*_1,70_ =4.74, *p*=.03, partial *η*^2^=.06, BF_10_=1.81, LHB amplitude *F*_1,70_=3.79, *p*=.06, partial *η*^2^=.05, BF_10_=1.22, Supplementary Fig. 1C). For additional LHB analyses, see Supplementary Results 5. Together the inferential and Bayesian statistics demonstrate that the CPP onset and build-up rate are robust age-related indicators.

**Extended Data Figure 3-3.**
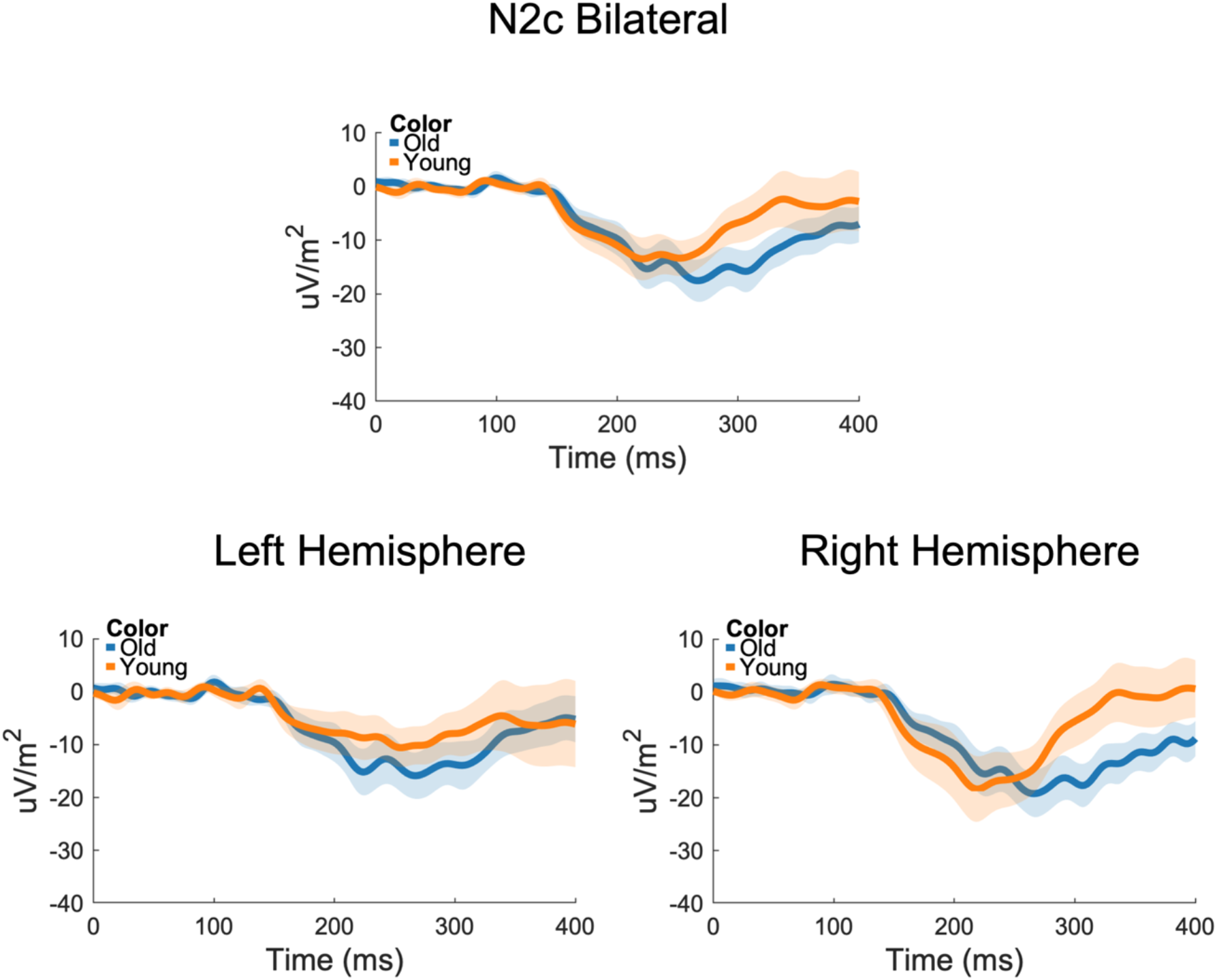
The N2c component, stimulus aligned at electrode P7 (Left Hemisphere), and P8 (Right hemisphere). Given the relevance of hemisphere lateralisation for theories of cognitive ageing, we investigated any age-related hemisphere differences in the N2c using 2 (old versus young) X 2 (right hemisphere x left hemisphere) ANOVAs, separately for latency and amplitude . There was no main effect of hemisphere on N2c latency (*F*_1,69_=3.46, *p*=.07) but there was a significant hemisphere x group interaction term (*F*_1,69_=11.56, *p*=.001, partial *η*^2^ =.14). In line with a large body of work highlighting a right hemisphere dominance for early sensory processing, follow up analyses revealed that the younger adults showed a significantly faster right hemisphere N2c latency (*M*=257.07ms, *SD*=51.67) as compared with the left hemisphere (298.77ms, 69.95; *F*_1,29_=14.99, *p*=.001, partial *η*^2^ =.34). In contrast, for the older adults there was no hemispheric differences in N2c latency (*F*_1,40_=1.23, *p*=.28; right hemisphere: *M*=303.59, *SD*=57.56; left hemisphere: *M*=291.37ms, *SD*=54.86), possibly indicative of a reduction in hemispheric asymmetries in the older adults. As compared with the younger adults, the older adults showed slower N2c latencies in the right (*F*_1,69_=12.32, *p*=.001, partial *η*^2^ =.15) but not left (*F*_1,69_=.25, *p*=.62) hemispheres. There was no effect of group on N2c amplitude (*F*_1,69_=3.33, *p*=.07), nor was there any group x hemisphere interaction term (*F*_1,69_=.02, *p*=.88).

**Extended Data Figure 3-4.**
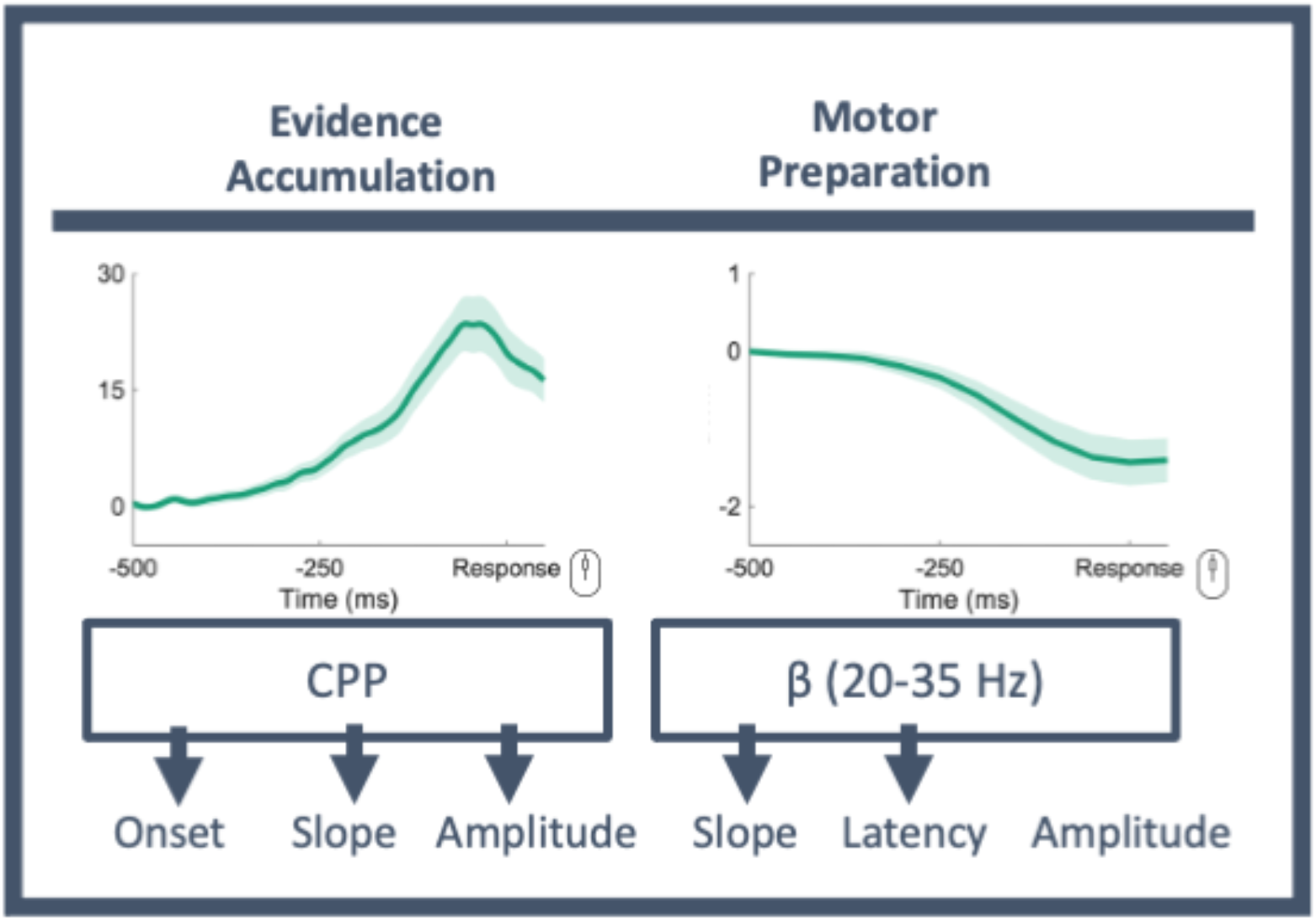
CPP and Beta components from Figure 1 visualised here aligned to participants’ response. To verify that motor preparatory activity was accurately captured by our stimulus-locked measure of beta latency, and to exclude the possibility that EE could be impacting RT through an influence over motor preparatory activity, two response-locked beta metrics were derived and explored in relation to RT: response-locked beta slope (build-up rate) and response-locked beta amplitude (threshold; Supplementary Fig. 4 below). Beta slope was defined as the slope of a straight line fitted to the response- locked waveform, with the time window defined individually for each participant between 300 to 50 ms pre-response, and baselined to -450 to -350ms. Beta amplitude was measured as the mean amplitude of a 100ms window centred on a participants’ response (i.e., -50 to +50ms around response). A stepwise linear regression model was used to identify which of the three beta measures (peak stimulus-locked latency, along with slope, and amplitude at the time of response) was the best predictor of RT (Criteria: probability of *F* to enter <=.05, probability of *F* to remove >=.1). The resulting model of RT included only stimulus-locked beta latency, indicating that this was the most appropriate EEG metric for capturing independent variance in RT (beta latency: standardized *β*=.54, *t*=5.25, *p*<.001, 95% CI [.28 .62]; beta slope: *β*=.10, *t*=.95, *p*=.34, beta amplitude *β*=-.12, *t*=-1.14, *p*=.26 model *F*1,68=27.60, *p*<.001). In line with previous work (e.g.(O’Connell et al., 2012; Brosnan et al., 2020), this result suggests that beta latency is a valid marker of task-relevant motor preparatory activity accounting for independent variance in response speed.

**Extended Data Figure 5-1.**
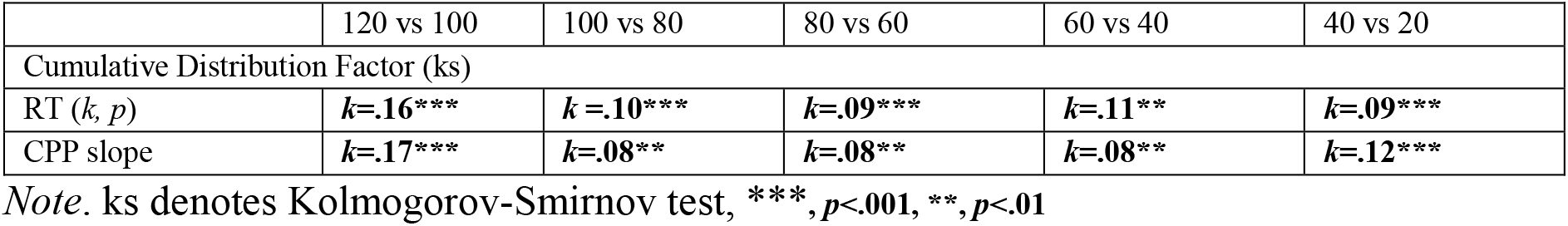

**Extended Data Figure 5-2.**
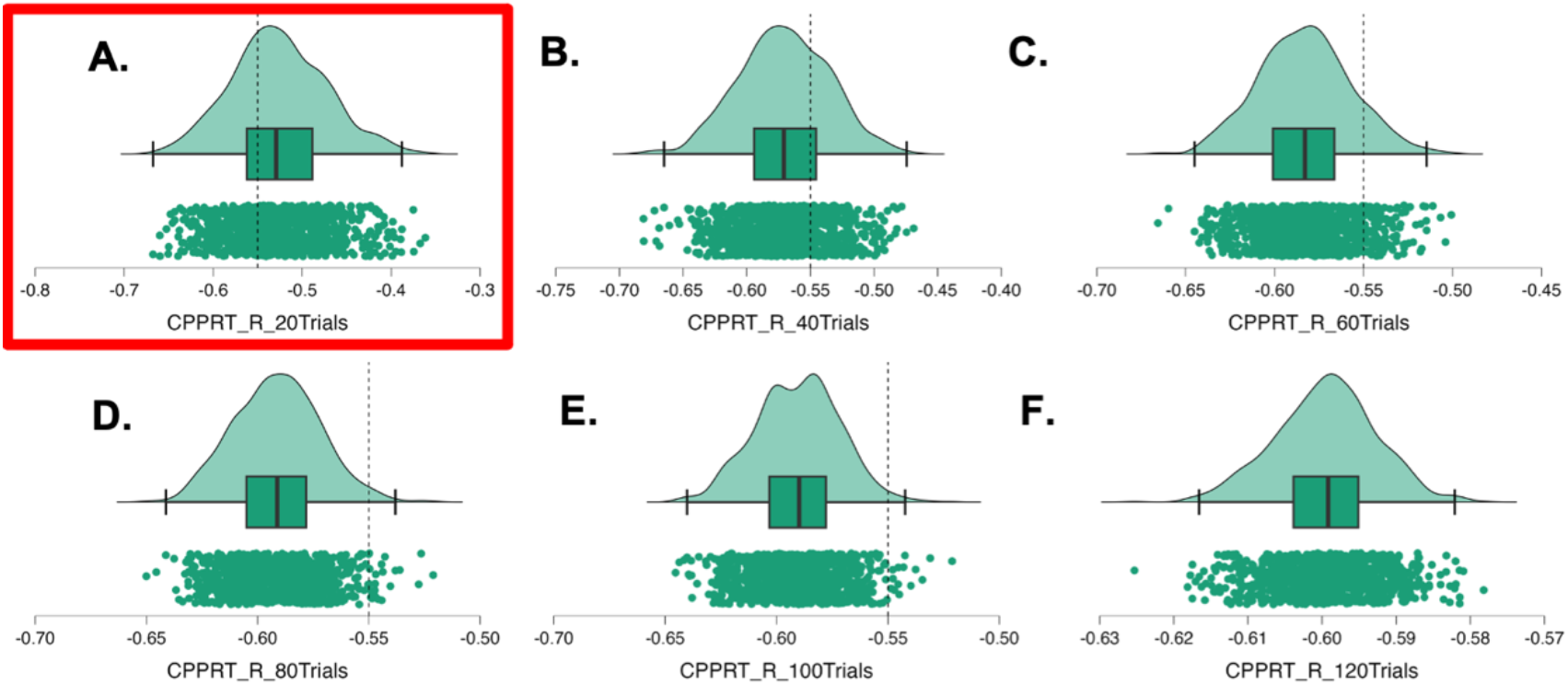
Each subplot denotes the direct relationship, Pearson’s *R*, between CPP build-up rate and RT for each of the 1000 permutations for each of the 6 bin sizes (20 up until 120 trials; A-F). Note the x-axis scales vary according to subplots. The dotted lines indicate a Pearson’s *r* value of -.55 between CPP build-up rates and Response Times (RT). Bayesian analyses indicate strong/infinite support for the alternative hypothesis that the effect sizes for the relationship between RT and CPP build-up rate with 120, 100, 80, 60, and 40 trials were larger than .55 (i.e, less than -.55 given the negative relationship between CPP build-up rate and response speed; all BF10>2.314ξ10^+63^). However, Bayes factor analyses revealed strong support for the null hypothesis (BF10=.002), i.e., that the estimates of effect size were not greater than -.55 (highlighted in red in **A.** above). **Calculation of the time necessary to assess 40 trials of the CPP**. Results from the minimum trial analyses indicate that a minimum of 40 trials would be sufficient to derive valid and behaviourally meaningful estimates of CPP build up rate. These trials are derived using the response-locked EEG signals, following data cleaning, for correctly identified target stimuli (coherently moving dots). Below we provide calculations both for the average time we expect necessary to obtain 40 valid trials, and for a ‘worst case scenario’. **Calculations using mean values**: Mean accuracy was 96%. In order to get 40 valid response locked trials, participants would need to be administered an extra 4% (2 trials), i.e., 42 trials in total. We calculated the percentage of EEG trials which were rejected by data cleaning i.e., (rejected trials/(rejected trials + valid trials), and on average 15% of trials were excluded following the preprocessing steps. In order to obtain 42 **valid** EEG trials (post data cleaning), an additional 15% of data would need to be collected, so 48 trials in total.

In this case, we would present the participant with

- 48 coherent motion trials at 3 seconds each

`o` = 144seconds total
- Each target trial would be preceded by random motion at three variable intervals (1.8s. 2.8s, & 3.8s; 48 targets/3 random motion periods =16 at each time period)
- [(1.8*16) +(2.8*16) + (3.8*16)]
- [28 + 44.8 + 60.8]

`o` =133.6seconds
- Total time (144s +133.6s) = 277.6 seconds (4.62 minutes)

### Calculations using the worst-case scenario values (i.e., participants with lowest accuracy and noisiest EEG data)

The participant with the lowest accuracy on the task correctly identified 71% of targets, and as such in order to get 40 valid trials would need an extra 29% (12 trials), i.e., 52 trials in total. While, on average 15% of participant EEG trials were rejected during processing, in the noisiest dataset we rejected 44% of trials. In this scenario, in order to obtain 52 valid trials post cleaning we would need an extra 44% of data (i.e., 75 trials in total).

In this case, we would present the participant with

- 75 coherent motion trials at 3 seconds each

`o` = 225 seconds total
- Each target trial would be preceded by random motion at three variable intervals (1.8s. 2.8s, & 3.8s; 75 targets/3 random motion periods =25 at each time period)
- [(1.8*25) +(2.8*25) + (3.8*25)]
- [45 + 70 + 95]

`o` =210seconds
- Total time (225s + 210s) = 435 seconds (7.25 minutes)

We note that these times do not account for EEG set-up time. Future work should address the minimum participant preparation time and reliability of these results from low density, portable electrode arrays.

**Extended Data Figure 6-1.**
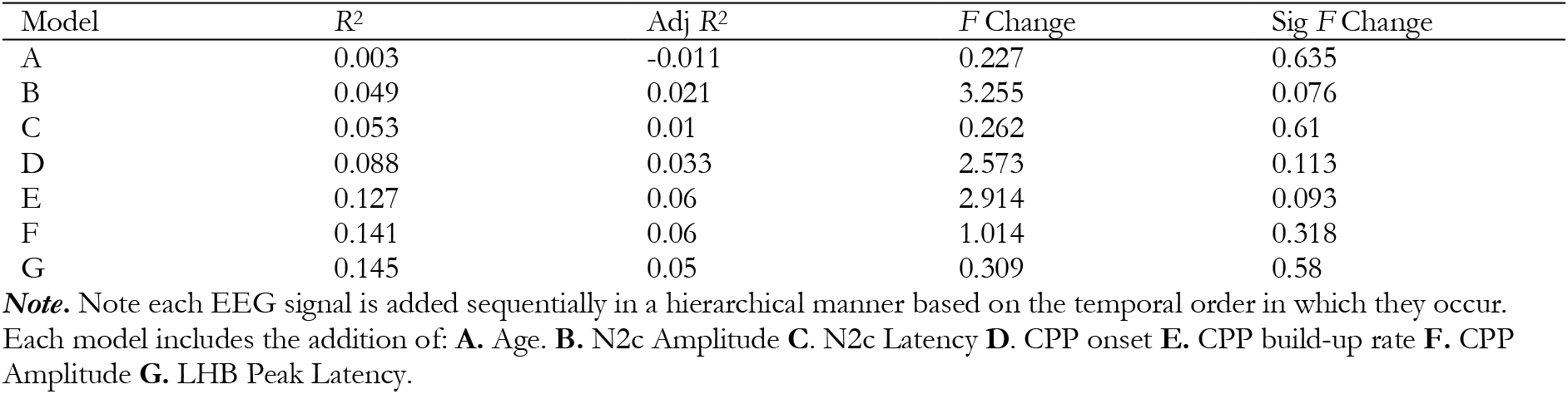
*t* parameter (non-decision time) modelled using a hierarchical linear regression model as a function of the EEG metrics.

**Extended Data Figure 6-2.**
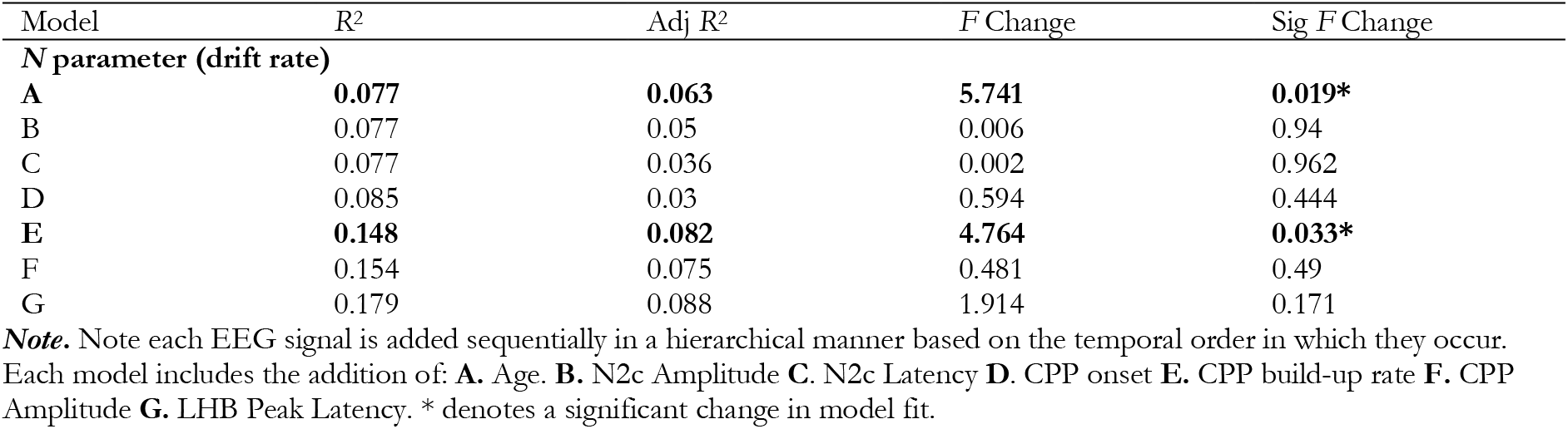
*ν* parameter (drift rate) modelled using a hierarchical linear regression model as a function of the EEG metrics.

**Extended Data Figure 6-3.**
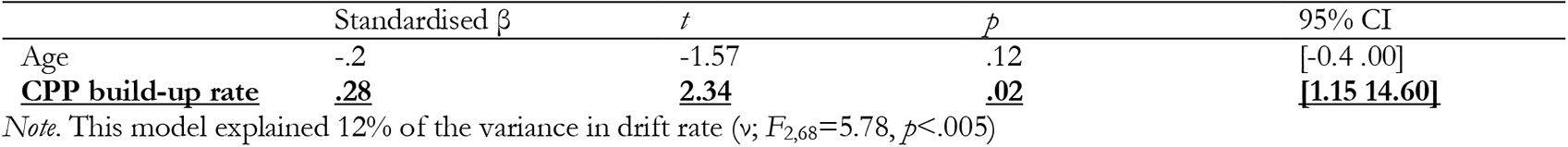
Final model of *ν* parameter (drift rate) as a function of the EEG metrics.

**Extended Data Figure 6-4.**
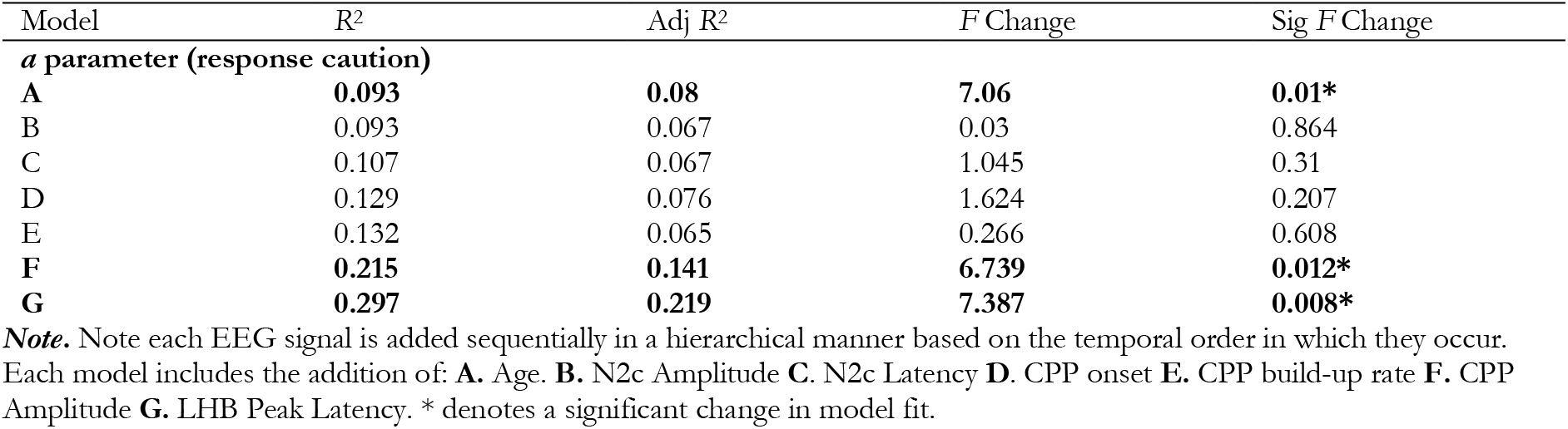
*a* parameter (response caution) modelled using a hierarchical linear regression model as a function of the EEG metrics.

**Extended Data Figure 6-5.**
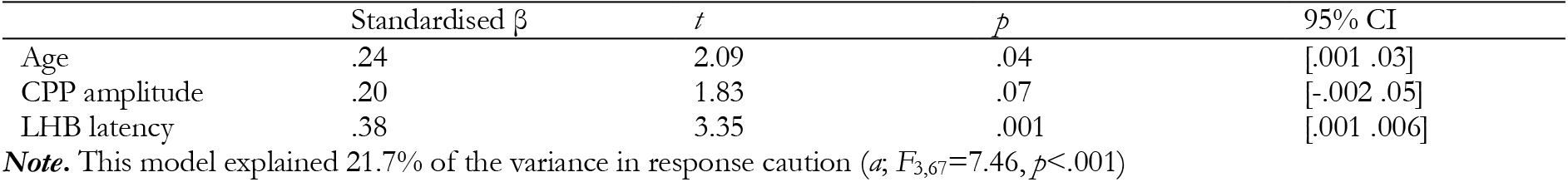
Final model of *a* parameter (response caution) as a function of the EEG metrics.

## ACKNOWLEDGMENTS

M.B.B. was supported by a Marie Skłodowska-Curie Fellowship from the European Commission (AGEING PLASTICITY; grant number 844246), and a Cullen Junior Research Fellowship at Corpus Christi College, University of Oxford. This work was additionally supported by the NIHR Oxford Health Biomedical Research Centre and the Wellcome Centre for Integrative Neuroimaging was supported by core funding from the Wellcome Trust (203139/Z/16/Z). M.A.B and R.G.O were supported by the Australian Research Council (DP150100986 and DP180102066). M.A.B. was further supported by a Senior Research Fellowship from the Australian National Health and Medical Research Council (APP1154378), and R.G.O by a European Research Council Consolidator Grant IndDecision 865474. ACN was supported by the Wellcome Trust (104571/Z/14/Z), a James S. McDonnell Foundation Understanding Human Cognition Collaborative Award (220020448), and an EU European Training Network grant (euSSN; 860563). T.C. was supported by the Australian Research Council (DP 180102383, DE 180100389), the Judith Jane Mason and Harold Stannett Williams Memorial Foundation, the Brain Foundation, and the Society for Mental Health Research. The funders had no role in study design, data collection and analysis, decision to publish or preparation of the manuscript. Finally, we thank Rory Boyle for helpful discussions on Cognitive Reserve. For the purpose of open access, the author has applied a CC BY public copyright licence to any Author Accepted Manuscript version arising from this submission.

